# Vascular smooth muscle cell atherosclerosis trajectories characterized at single cell resolution identify causal transcriptomic and epigenomic mechanisms of disease risk

**DOI:** 10.1101/2025.06.04.655863

**Authors:** Daniel Yuhang Li, Soumya Kundu, Paul Cheng, Wenduo Gu, Will Jackson, Quanyi Zhao, Trieu Nguyen, Matthew Worssam, Joao Pinho Monteiro, Roxanne Diaz Caceres, Stanley Dale, Brian Palmisano, Chad S Weldy, Ramendra Kundu, Anshul Kundaje, Robert Wirka, Thomas Quertermous

**Affiliations:** Division of Cardiovascular Medicine, 300 Pasteur Drive, Falk CVRC, Stanford, CA; Department of Genetics, Stanford University School of Medicine, Stanford, CA; Univ. of North Carolina, Dept. of Medicine, Chapel Hill, NC, USA

## Abstract

Vascular smooth muscle cells (SMC) contribute to heritable coronary artery disease (CAD) risk and undergo complex cell state transitions to multiple disease related phenotypes. To investigate the genetic basis of SMC state trajectories that underlie the SMC component of CAD causality we have developed a dense timecourse single cell transcriptomic and epigenetic map of atherosclerosis in a murine disease animal model. Cellular trajectories were derived from the temporal data and probabilistic fate modeling with Waddington-Optimal Transport (WOT). We created transcription factor (TF) centered regulons mapped across the developmental timeline and through network-based prioritization with WOT predicted TFs and in silico TF perturbation, identified key drivers of cell state changes associated with EMT, vascular development, and circadian clock functions. Integration of mouse disease data with human CAD genetic findings identified transition SMC phenotypes that mediate disease risk and point to causal disease mechanisms. Parallel studies using knockout of the validated CAD gene Tcf21 revealed its impact on SMC transition cellular phenotypes and disease risk genes, due in part to a role regulating the transition of SMC precursor cells in the secondary heart field. Together, these studies characterize atherosclerosis trajectories at single cell resolution and identify genetic causal transcriptomic and epigenomic mechanisms of CAD risk.

## Introduction

Cardiovascular diseases, principally coronary artery disease (CAD) and stroke, are the worldwide leading cause of global mortality^1^. Therapies that modify classical environmental and metabolic factors have ameliorated a portion of CAD risk^2^, but more than half of the risk can be attributed to common inherited genetic variation by affecting vessel wall pathways that mediate disease pathophysiology. These genetic factors remain unidentified and untreated^3–6^. While extensive studies have investigated the cellular and molecular features of atherosclerosis, they have been unable to establish causality and thus hinder the translation towards vascular wall directed therapies^7^.

The smooth muscle cell (SMC) lineage, which appears to make the largest contribution to disease risk^8–10^, undergoes extensive complex phenotypic transitions that have not been well characterized at the cellular and molecular level, making it difficult to assign causality to specific gene programs^11–16^. The few SMC GWAS genes studied in detail suggest that complex transcriptional programs can direct cellular trajectories that lead to fibroblast-like (fibromyocyte, FMC) or osteochondrogenic (chondromyocyte, CMC) phenotypes, with these cellular phenotypes being proposed to mediate opposing effects on disease risk^12,13,17^. However, this paradigm is in conflict with genomic analyses of limited single cell data suggesting that all FMC transition to CMC over the course of atherosclerotic lesion development^9,12,18^. Molecular and cellular studies of SMC transitions to human genetic data linking locus associations to individual SMC gene causality and direction of effect, are needed to promote progress in the field.

Studies reported here are aimed at the comprehensive characterization of genes and gene programs that function in SMC to mediate phenotypic transitions and disease risk. We describe a mouse transcriptomic and epigenomic atlas of atherosclerosis with single cell RNA and chromatin accessibility (ATAC) data collected over a dense timecourse in a well-accepted mouse model. For the first time, we map the SMC lineage cellular trajectories with real-time course gene expression, chromatin accessibility data and advanced trajectory inference methods such as the Waddington Optimal Transport algorithm, identifying the genes and gene regulatory networks that mediate the transitions to FMC and CMC cellular phenotypes. Further, we investigate how these trajectories are affected by the CAD protective *Tcf21* gene, mapping the regulatory network downstream of this gene, further identifying collaborating transcription factors that mediate genome-wide regulation of SMC phenotype transitions and disease risk.

### Construction of an SMC lineage traced multi-omic single cell timecourse atherosclerosis cell atlas

We performed single-cell RNA sequencing (scRNAseq) and single-nucleus assay of transposase-accessible sequencing (scATACseq) with aortic root tissue from *ApoE-/-* (*ApoE* KO) mice to characterize the complex genetic regulatory networks that control the developmental cascade of smooth muscle cell (SMC) phenotypic transitions in the context of atherosclerotic stress^13^. These mice, also expressing a tamoxifen-inducible Cre recombinase driven by the SMC-specific *Myh11* promoter and a Cre-activatable *tdTomato* reporter gene, were fed a high fat diet (HFD) across 7 timepoints for scRNAseq and 6 timepoints for scATACseq assays **(Fig. 1A)**. Aortic root tissues were collected, digested, and subjected to droplet-based cell capture, and independent RNA and transposed DNA sequencing^13^. The resulting data were subjected to dimensionality reduction, clustering and visualization with Seurat, providing individual cell clusters that we and others have identified previously **(Fig. 1B)**^11–16^. For this study, we subsetted lineage traced SMC clusters **(Fig. 1C, S-Fig. 1A-H, S-Table 1, Methods)**, transferred scRNASeq labels onto scATAC cells and co-embedded RNA and ATAC modalities, resulting in a total of 96,195 high quality cells across both modalities (**Figs. 1C, D)**^13,19^.

**Figure 1.**
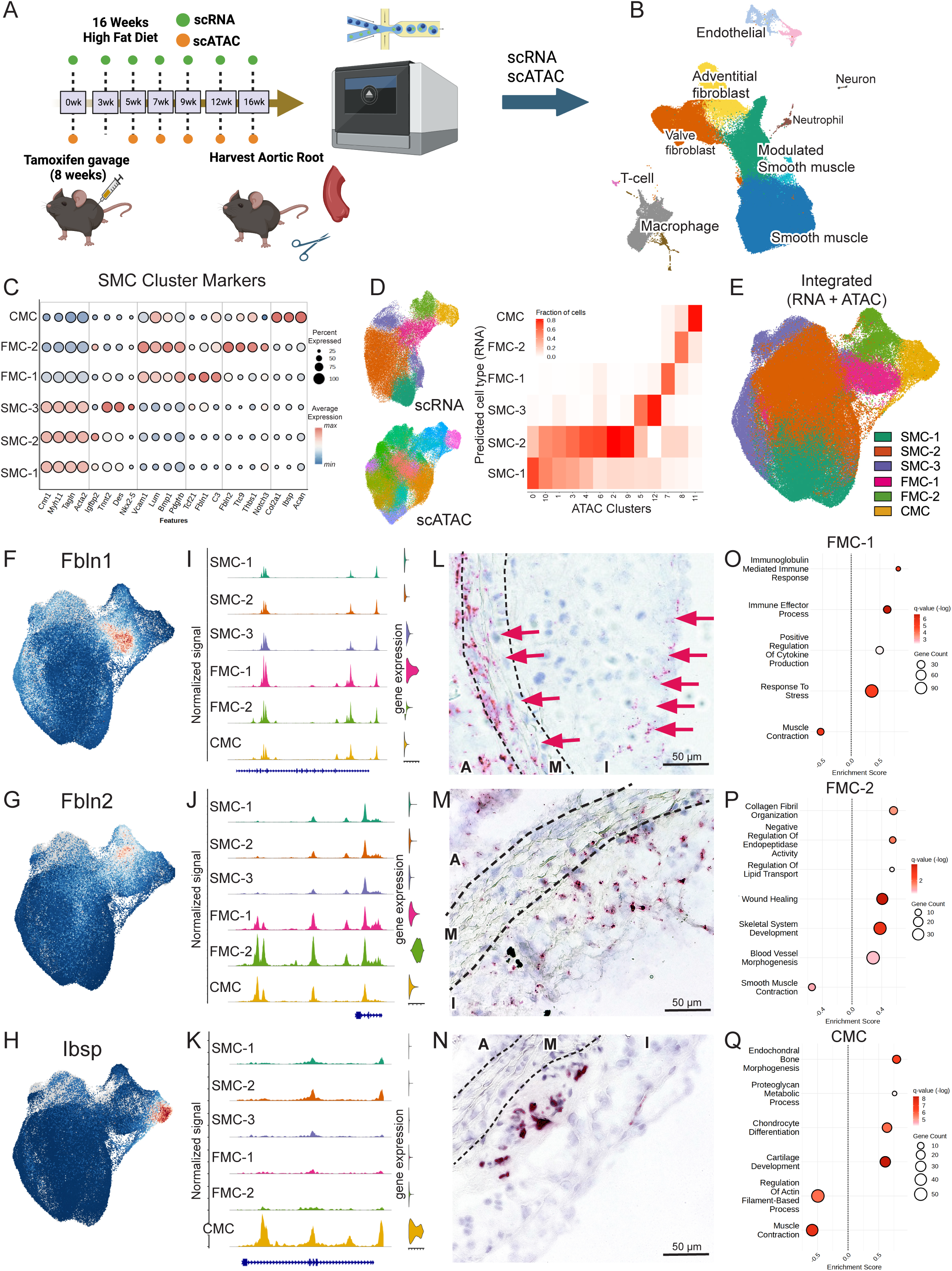
Mouse atherosclerosis timecourse model and SMC cluster identity. **A.** Schematic of the single cell scRNA and scATAC sequencing data collection in the *ApoE* KO mouse atherosclerosis model. Mice were tamoxifen gavaged at 8 weeks of age to induce *Myh11-Cre* recombination and *tdTomato* expression, single cells are isolated by flow cytometry and sorted on lineage traced and non-lineage traced cells. Mouse aortic root tissues were collected and scRNAseq assays performed on single cells isolates after 0, 3, 5, 7, 9, 12 or 16 weeks of high fat diet feeding. Remaining single cells from the same cellular samples underwent nuclei isolation and scATACseq assays at the 0, 5, 7, 9, 12, and 16 week timepoints. **B.** UMAP embedding of all aortic root resident cells, with scRNAseq cluster assignment guided by cell-specific gene expression as we have described ^19^. **C.** Dotplot identifying selected top markers on a per-cluster basis (expanded in S-Fig. 1). **D.** UMAP of scRNA and scATAC clusters prior to integration. Confusion matrix showed cluster-cluster mapping between scRNA (x-axis) and scATAC (y-axis) cluster cell assignment data after CCA co-embedding. **E.** UMAP visualization of the integrated object with scRNA and scATAC co-embedding. **F-H.** Feature plots for specific marker genes *Fbln1* (FMC-1), *Fbln-2* (FMC-2), and *Ibsp* (CMC) in SMC. **I-K.** Coverage plots of scATACseq open chromatin peaks for marker genes *Fbln1*, *Fbln2*, and *Ibsp*. **L-N.** In situ RNA hybridization (RNAScope) at 20x magnification for respective marker genes indicated that the three transition phenotypes were differentially localized in the vessel wall. (A – Adventitia, M – Media, I – Intima, scale bar – 50 µm) **O-Q.** GSEA enrichment of cluster specific gene ontology biological process pathways for each modulated SMC cluster.

### Distinct molecular and spatial patterns characterize SMC-derived lineage cells

We expanded upon previously identified SMC cell states^11–16,19^ and identified six distinct SMC lineage phenotypes supported by both RNA and chromatin accessibility patterns **(Fig. 1E, S-Fig. 1I, J, S-Table 2)**. We observed high level expression of canonical SMC differentiation markers (*Cnn1*, *Tagln*, *Acta2*, *Myh11*) in three clusters (SMC-1, 2, 3). While SMC-1 primarily expressed these markers, SMC-2 demonstrated expression of markers such as *Igfbp2*, indicating an early phenotypic transition state^20,21^. A distinct SMC-3 population additionally enriched for myocardial markers *Tnnt2* and *Nkx2-5*, identified the previously characterized population of SMC at the base of the aortic root arising from the SHF^21,22^.

All FMC were characterized by robust expression of fibroblast and epithelial mesenchymal transition (EMT) markers such as *Vcam1*, *Lum*, *Bmp1 and Pdgfrb* **(Fig. 1C. S-Fig. 1K)**. However, we identified two FMC sub-populations demonstrating discrete patterns of gene expression **(Fig. 1F,G)**, chromatin accessibility **(Fig. 1I, J)**, spatial localization **(Fig. 1L, M)**, and functional enrichment **(Fig. 1O, P)**. ATAC data showed overlapping but distinct regions of open chromatin, solidifying the integrity of each FMC molecular subtype **(Fig. 1D, I, J)**. One group termed FMC-1, expressed specific markers including *Tcf21, Fbln1,* and targets of interferon gamma signaling including *H2-Aa/Ab1/Eb1*, *Cd74* and *Il33* (**S-Fig. 1K)**. In situ RNA hybridization with cluster-specific FMC-1 markers demonstrated expression largely encompassing the media and fibrous cap **(Fig. 1L, S-Fig. 2A-C)**. Transcriptional functional enrichment with gene set enrichment analysis (GSEA) using GO biological process gene sets for FMC-1 cells identified processes such as “immune activation” and “cytokine production” **(Fig. 1O, S-Table 3).** The second cluster of cells, termed FMC-2, were identified with specific markers *Thbs1* and *Notch3* localized to the fibrous cap **(S-Fig. 2D, E)** or spanning the intimal plaque (*Col8a1, Fbln2, Ttc9*) **(Fig. 1M, S-Fig. 2F, G, H)**. Although FMC-2 shared a number of terms with FMC-1, expressed genes were uniquely associated with “collagen fibril organization”, “regulation of lipid transport” and “wound healing” **(Fig. 1P).** Interestingly, a number of genes (*Loxl1, Bmp1, Vcam1*) were found to have vascular wall expression patterns encompassing those seen for both FMC-1 and FMC-2, possibly reflecting cells in transition **(S-Fig. 2I, J, K).**

By comparison, the CMC cluster identified a distinct transition with a highly restricted chromatin accessibility pattern, and restricted expression of marker genes *Col2a1* and *Ibsp* localized to the base of the plaque **(Fig. 1N, S-Fig. 2L, M)** as we have shown previously^13,17,19^. CMC pathways were consistent with osteochondrogenic processes, as described by several groups^11–16,19^ **(Fig. 1Q, S-Table 3)**.

To translate these spatial findings from mice to human, we further performed label transfer onto publicly available scRNA data from human donor transplant coronary arteries and observed a similar distribution of FMC-1, FMC-2 and CMC by single cell features and spatial transcriptomics^23^ **(S-Fig. 2N-S)**.

### A probabilistic model of SMC atherosclerosis trajectories reveals distinct patterns of gene expression during SMC cell state transitions

We utilized force directed layouts to visualize transcriptional relationships, in real-time, across the SMC atherosclerosis trajectories **(Fig. 2A)**. We first observed the appearance of FMC-1 cells at 5 weeks of HFD with a subsequent increase in FMC-2. This observation is consistent with a migratory path for SMC lineage cells from the media to the fibrous cap and subsequently down into the intimal plaque^24–28^ **(Fig. 1L, M**; **Fig. 2A, B, C).** Consistent with this possibility, FMC-2 were also enriched for osteoblast progenitor gene signatures^29^ **(S-Fig. 3A)**. CMC abundance dramatically increased at 12 weeks of high fat diet, correlating with spatial localization at the plaque base **(Fig. 1N)**. In addition, transcriptomic patterns suggested increasing pathologic signatures including senescence, EMT, angiogenesis, apoptosis, and efferocytosis **(S-Fig. 3B-F)** as the cells transitioned towards the CMC cell state at the plaque base, which is consistent with the acellularity in this region^13,25^.

**Figure 2.**
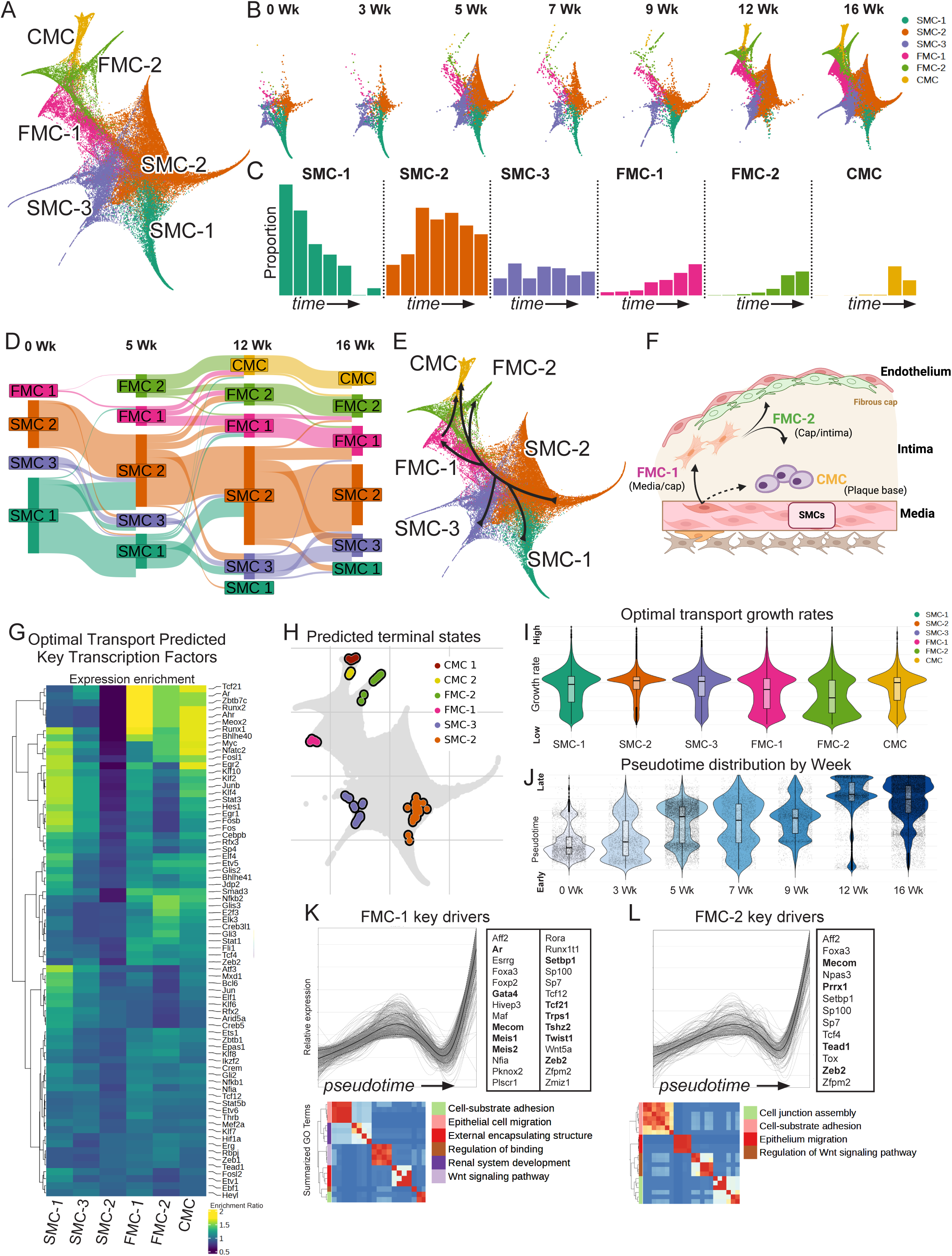
Characterization of disease related SMC lineage transition cell trajectories. **A.** Force directed layout (FLE) representation of lineage traced SMC showing clustered cell phenotypes. **B.** FLE embedding of lineage traced SMC split by weeks on high fat diet. **C.** Cluster proportions of lineage traced cells split by week. **D.** Sankey plot of Waddington optimal transport (WOT) predicted transition probability from SMC to FMC and CMC populations. Band thickness indicates relative transition probability from starting to end cell states. **E.** FLE embedding of SMC lineage transition cell states with arrows representing WOT predicted movement of cells during phenotypic transition. **F.** Illustrated representation of WOT predicted movement of cells during phenotypic transition. **G.** Transcription factor enrichment for cells fated to become each cluster-type calculated by WOT based on week 5 to week 12 transition. **H.** CellRank2 derived predicted terminal cell states. **I.** Optimal transport derived growth rates by cell state. **J.** Pseudotime distribution of cells by week. **K-L.** CellRank2 summarized expression trends across pseudotime (right) showing a representative activating gene cluster towards FMC-1 or FMC-2, TFs within this gene cluster (middle), and GO enrichment of cluster genes (left) for key genes.

To quantitatively model these observed cell state changes, we applied the Waddington Optimal Transport (WOT) algorithm to build a probabilistic model for cell state transitions^30,31^. WOT is a heuristic method that models growth rates based on cell cycle and apoptosis gene expression to perform developmental trajectory inference. Applying this algorithm across timepoints, we found that while SMC-1/2/3 are present at baseline (Week 0), SMC-2 are the primary cell phenotype that transitions to FMC states **(Fig. 2D, E, F)**. WOT modeling suggested that a small FMC-1 population existed at baseline, significantly expanded by 5 weeks, and directly contributes to both FMC-2 and CMC. FMC-2 appeared to primarily become CMC which were evident by 12 weeks of HFD. However, given the higher proportion of FMC-1 present relative to FMC-2, both FMC cell states likely contributed similarly to the total number of CMC. Further, it is important to note that FMC-1 and FMC-2 both show an initial proportional increase that is sustained at the 16 week timepoint, suggesting that their phenotypes may both be terminal endpoints for SMC transition^32^. This is in contrast to previous pseudotime based analysis which have suggested that CMC are the common primary endpoint for all transitioning SMC in the plaque^9,12^.

WOT further captured TF enrichment across cell state transitions, using weeks 5 and 12 as reference points which encapsulated the majority of the cell state transitions. This approach allowed us to highlight the crucial TFs expressed in SMC that guide these cells towards their final fates **(S-Table 4)**. We then filtered these transition TFs with significantly enriched accessible transcription factor binding motifs identified from scATACseq data. This integrative approach enabled us to identify TFs that have previously been associated with SMC phenotypic transitions, and in some cases pointed to novel functions for these genes **(Fig. 2G)**. For example, we found *Tcf21* expression to be the most enriched TF in cells fated to become FMC-1, consistent with its early role in phenotypic transitions, while WOT-based TF enrichment also suggested a prominent role in the FMC-CMC state transition. Other genes that inhibit cellular proliferation, *Meox2*^33^ and *Zbtb7c*^34^, showed activation of expression in FMC-1 development, persisting into cells in the CMC state, suggesting a role for precise regulation of cell expansion throughout the phenotypic transition process. Well studied genes such as *Runx1* and *Runx2* were predicted to mediate the FMC-CMC transition. However, enriched expression of these genes in FMC-1 and FMC-2 phenotype fated cells is surprising and suggests that they may also have a role promoting suppression of the SMC phenotype. Indeed, like *Tcf21*, *Runx2* has been shown to block transactivation of the *Myocd-Srf* program by interacting directly with *Myocd* to inhibit binding to the CArG box in SMC genes^35,36^. Other genes known to promote chondrogenesis such as *Nfatc2,* which also promotes SMC lineage proliferation^37–39^ were noted to be enriched in the CMC state. Moreover, the top SMC-3 promoting TF was *Isl1*, reinforcing its SHF origins^40^ **(S-Table 4).**

The suggested trajectory paradigm is further supported by analyses with the RealTime kernel calculations in CellRank2 that computationally derive terminal states. This analysis recovered FMC-1, FMC-2, two CMC and the SMC-2 and SMC-3 as terminal states **(Fig. 2H**). Further, we visualized the WOT derived growth rates by cell state and found a greater proportion of cells with increased proliferation in the early transitioning SMC-2 cells, reduced growth rates in the FMC with FMC-2 having higher proportion of low growth rates and higher senescence score **(Fig. 2I, S-Fig. 3B),** and lastly a higher growth rate in CMC, reflective of proliferative early-stage chondrocytes in developing bone^41,42^ **(Fig. 2I).** Given the high correlation between pseudotime and cell state stages confirmed by our real-time analysis (pearson’s r = 0.57; spearman’s rho = 0.56, p < 2e-16) **(Fig. 2J)**, we modeled the atherosclerosis trajectory with CellRank2 as an orthogonal approach to derive a probabilistic transition model. The increased resolution provided by pseudotime allowed us to cluster gene expression across the different inferred trajectories, identify key TFs within trajectory gene clusters, and characterize pathways based on gene expression trends clustered over pseudotime. For example, we found that FMC-1 trajectory clustered genes demonstrate early activation of genes involved in “inflammatory response” and “response to molecules of bacterial origin” suggestive of a core set of genes involved in stress response including TFs such as *Klf4*, *Cebpb*, *Runx1* and *Nfkb1* **(S-Fig. 4, cluster 3)** followed by gene groups demonstrating “epithelial cell migration” and “cell-substrate adhesion” including TFs like *Tcf21*, *Ar*, *Zeb2*, and *Twist1* **(Fig 2K)** and more unique processes such as “antigen processing and presentation” and “cytokine mediated signaling” that includes TFs such as *Ahr*, *Gas7* and *Stat1* **(S-Fig. 4, cluster 2).** For FMC-2, we observed an early cluster enriched for “epithelium migration” and “regulation of Wnt signaling” **(S-Fig. 5, cluster 3)**, while later clusters showed enrichment for “cell-junction assembly”, “cell chemotaxis”, “angiogenesis” and “collagen metabolic process” **(Fig 2L, S-Fig. 5, clusters 1, 2).** Similarly, for CMC, there were robust signals for “biomineralization” and “ossification” terms **(S-Fig. 6).**

### ATACseq data identifies cell state specific transcription factor motifs

To characterize the epigenetic processes that mediate the noted transcriptional effects, we investigated chromatin accessibility in the transitioning cells with scATACseq data **(Fig. 1D).** We observed high specificity of chromatin accessibility within each label transferred *tdTomato* lineage traced cluster and visualized top specific peaks along pseudotime **(Fig. 3A)**. Using GREAT for functional enrichment^43^, we identified specific cellular processes including smooth muscle related processes in the SMC, “cell migration”, “inflammation” and “response to platelet derived growth factor” in the FMC, and “ossification” and “chondrocyte development” in the CMC **(Fig. 3A, S-Table 5).**

**Figure 3.**
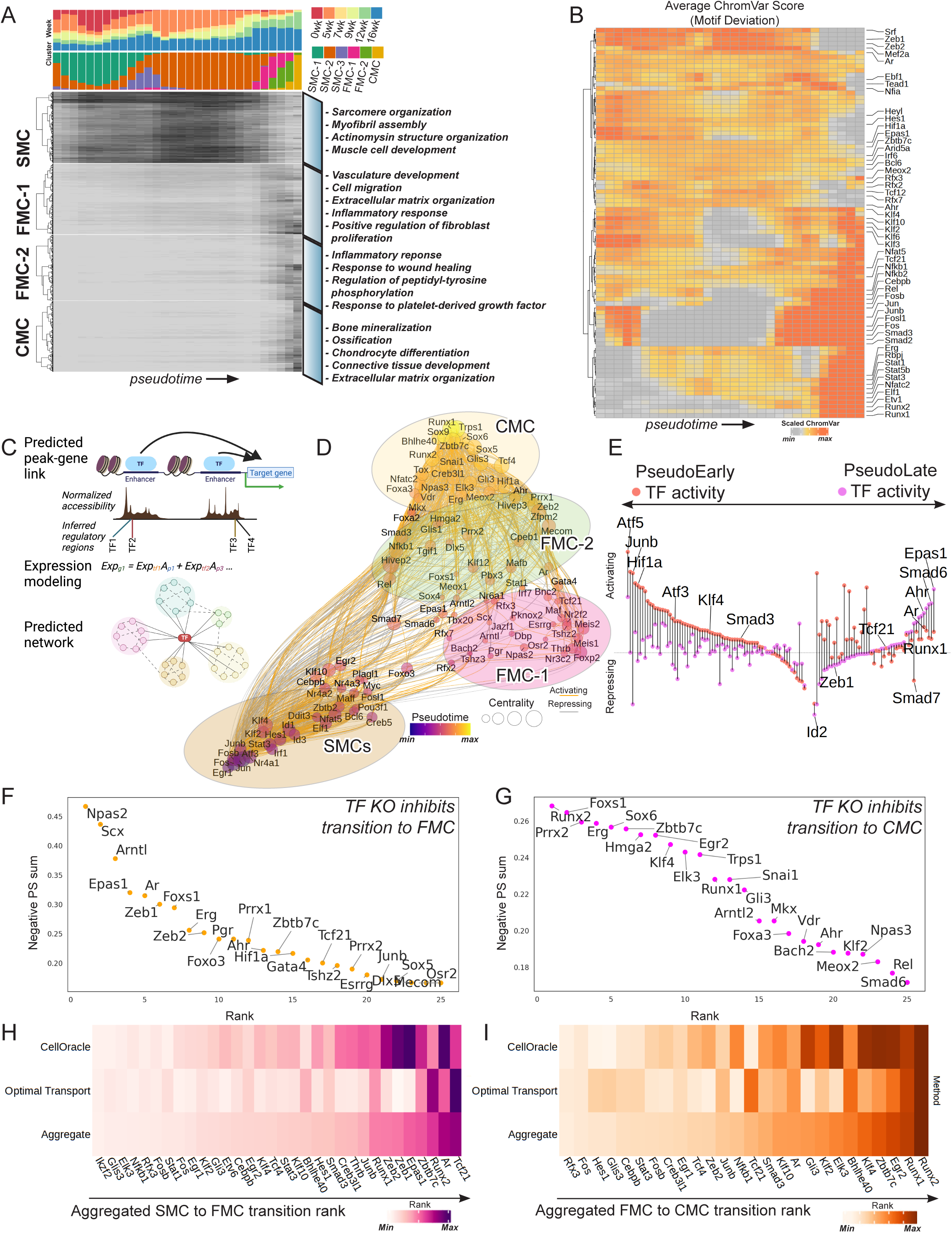
Integrative analysis identifies core accessible transcription factor motifs and provides network inferences. **A.** Hierarchical clustering of top specific scATAC peaks arranged by cell states across 30 pseudotemporal bins. Top two bars show cell proportion by week and by cluster assignment, respectively. Heatmap displays min-max scaled mean chromatin accessibility for each SMC cell state. Representative biological processes from GREAT pathway enrichment for each cell state is displayed on the right. **B.** Hierarchical clustering of ChromVar scores (motif accessibility deviation from background) for cluster enriched motifs across 30 pseudotemporal bins. **C.** Representative schematic of network inference. TF binding through binding site-motif matching within accessible chromatin regions are linked to genes. The expression of TF target genes were then fitted with a bagging regression model to generate TF-target gene networks. **D.** Summarized PANDO predicted TF network represented with UMAP embedding. Color gradient and size represent the expression-weighted pseudotime and centrality scores, respectively. **E.** Differential TF activity between SMC (salmon) and modulated SMC states (purple). TF activity is calculated by multiplying the mean regulatory coefficient of each TF network with its respective average TF expression. Sign of the activity represents inferred net regulatory effect – activating (+) or repressing (-). **F,G.** Ranked perturbation scores (product between control differentiation and KO-simulation vectors) for systematic KO simulation effect of TFs on the SMC and FMC lineages. Higher scores predict decreased likelihood of phenotypic shift from the originating cell state after TF perturbation. **H-I.** Rank of top TFs predicted to affect the SMC to FMC or FMC to CMC transition using 1) CellOracle in silico perturbation scores, 2) Optimal Transport derived fated cell TF enrichment, or 3) aggregated ranking across both methods.

To prioritize dynamic TF binding motifs, we applied ChromVar to calculate their motif binding accessibility probability distributions. We filtered these data using a core set of overrepresented TF motifs identified by Signac and HOMER and visualized motif deviations along pseudotime to capture the dynamic shifts of TF binding activity **(Fig. 3B)**. We used the early pseudotime bins 3-24 (PseudoEarly) and late pseudotime bins 25-30 (PseudoLate) to approximate the SMC-FMC and FMC-CMC transitions (**Fig. 3A**). These analyses identified temporal trends for both known and novel regulators of smooth muscle fate. For example, Srf, Mef2, and Zeb motif deviation was observed to be higher in SMC^13,44^. In addition, for the early transitions, we observed a number of TFs with the greatest motif accessibility across the SMC stages, including Tead, Zbtb7c, Meox2, Rfx, and Nfi factors. Factors showing greatest motif accessibility during the late transition stages included Tcf21, Smad3, Rbpj, Stat, Nfatc, Cebpb, and Runx factors, with many of these known to be involved in endochondral bone formation. KLF factors and numerous AP-1 factors showed a bimodal pattern with accessibility early in the quiescent SMC state and later, from FMC to pre-CMC bins, likely representing their pioneer factor functions at these differentiated cell states. These analyses summarized the chromatin landscape guiding TF pathways, extending previous studies using pseudotime data derived from baseline and terminal phenotypic cell states^13^ **(Fig. 3B)**.

### Integration of network-based TF prioritization identifies *Tcf21* as a top candidate to alter SMC to FMC transition

Using our co-embedded scRNA and scATAC dataset, we leveraged complementary network inference methods, Pando^45^ and CellOracle^46^, to create a custom workflow to infer transcription factor-target interaction networks (gene regulatory networks, GRNs) that direct phenotypic transition and simulate cell identity changes with *in silico* TF perturbations **(Fig. 3C, S-Tables 6,7, Methods)**. The dataset was divided by PseudoEarly and PseudoLate bins to infer GRNs for these analyses. We visualized a summary GRN colored by the average pseudotime-TF expression to reveal a cascade of network activation **(Fig. 3D)**. SMC lineage phenotypes clustered separately, identifying numerous TFs that are likely to direct specific regulons that mediate cell states and transitions characteristic of the response to disease stresses **(S-Table 8).** We computed TF activities for the PseudoEarly and PseudoLate SMC states, and a comparison of TF activity change from SMC to transitioning SMC cell state activity revealed novel patterns such as shifts in hypoxia inducible factor (HIF) activity highlighted by the known interaction between *Hif1a* and *Epas1*^47^ and other factors such as *Ahr* that interact with HIF through the common heterodimer *Arnt*^48^ **(Fig. 3E)**.

We then performed systematic *in silico* perturbations using our inferred TF-target links from SMC-FMC and FMC-CMC transitions to calculate perturbation scores measuring the potential of TFs to drive cell transition away from the originating cell state. In SMC, we identified enrichment of EMT factors such as *Tcf21* and *Zeb* factors, and similar to the top TF activity changes, we identified enrichment for hypoxia inducible factors including *Hif1a*, and *Epas1*, reflecting their high network connectivity in early phenotypic transitions **(Fig. 3F).** For factors promoting change to CMC phenotypes, we identified known factors such as *Klf4*^11^ and *Runx1/2*^49,50^, in addition to unique factors such as *Trps1, Sox6, Erg, Zbtb7c, Prrx1, Arntl2*, and *Snai1* that were nominated to drive the differentiation of cells from FMC to CMC **(Fig. 3G)**.

Lastly, we created an aggregated TF transition ranking by combining normalized enrichment from WOT for core TFs promoting the FMC-1 and CMC fates and normalized SMC-FMC as well as FMC-CMC perturbation scores from CellOracle (unfiltered WOT and CellOracle heatmaps in **S-Fig. 7A-C**). Through this combination of cell fate enrichment and network-based prioritization analyses, we found *Ar, Zeb, Smad, Mecom, Prrx1, and Cebpb* factors*, as well as Tcf21*, to be among central enriched TFs involved in the phenotypic transition from SMC to FMC **(Fig. 3H, S-Fig. 7B)**, and *Runx, Klf, Egr2, Sox9, and Zbtb7c* as top drivers for FMC to CMC transition **(Fig. 3I, S-Fig. 7C**). Additionally, *Runx* and *Zbtb7c* factors were consistently highly ranked across both transitions.

### Loss of *Tcf21* expression results in decreased FMC and CMC cell numbers, and altered cell state transition probabilities

As a top TF predicted to direct SMC to FMC transition, we further examined the effect of exemplar CAD gene *Tcf21* knockout on SMC trajectories with updated single cell chemistries using the timecourse methods employed for the control dataset to elucidate *Tcf21* SMC regulatory mechanisms that occur with vascular stress.

We first examined changes in SMC transition cell numbers that resulted from *Tcf21*-KO. In keeping with our prior work^16^, there was a significant decrease in the FMC populations at 12 and 16 weeks of HFD **(Fig. 4A)**. Comparison of lineage-traced SMC proportions revealed a marked ∼3-fold increase in the SMC-3 *Tcf21*- KO cells at 5 weeks that persisted to week 12. These *Tnnt2*-expressing cells have been lineage traced to the secondary heart field^21,22^, suggesting an early expansion of this aortic base contractile medial compartment in the context of *Tcf21* loss. The possible involvement of *Tcf21* in the regulation of these cells was supported by its expression in *Nkx2-5* lineage traced cells as identified with scRNASeq **(S-Fig. 8A-E)**. At 12 weeks, there was a notable relative decrease in *Tcf21*-KO FMC-1, FMC-2 and CMC proportions that corresponded with a relative increase in SMC-2 and SMC-3 cluster proportions, suggesting that the *Tcf21*-KO cells were halted in disease associated transitions. At 16 weeks, there was a consistent decrease in FMC-1 without a decrease in FMC-2, suggesting potential compensation for the loss of *Tcf21* expression in the SMC lineage cells over time **(Fig. 4A)**. At both 12 and 16 weeks we also observed a decrease in CMC cell proportion in the *Tcf21-*KO relative to control, suggesting that decreased FMC leads to decreased CMC via a mass action effect, or that *Tcf21* directly promotes CMC development as suggested by previous analyses **(Figs. 2G, 3I).**

**Figure 4.**
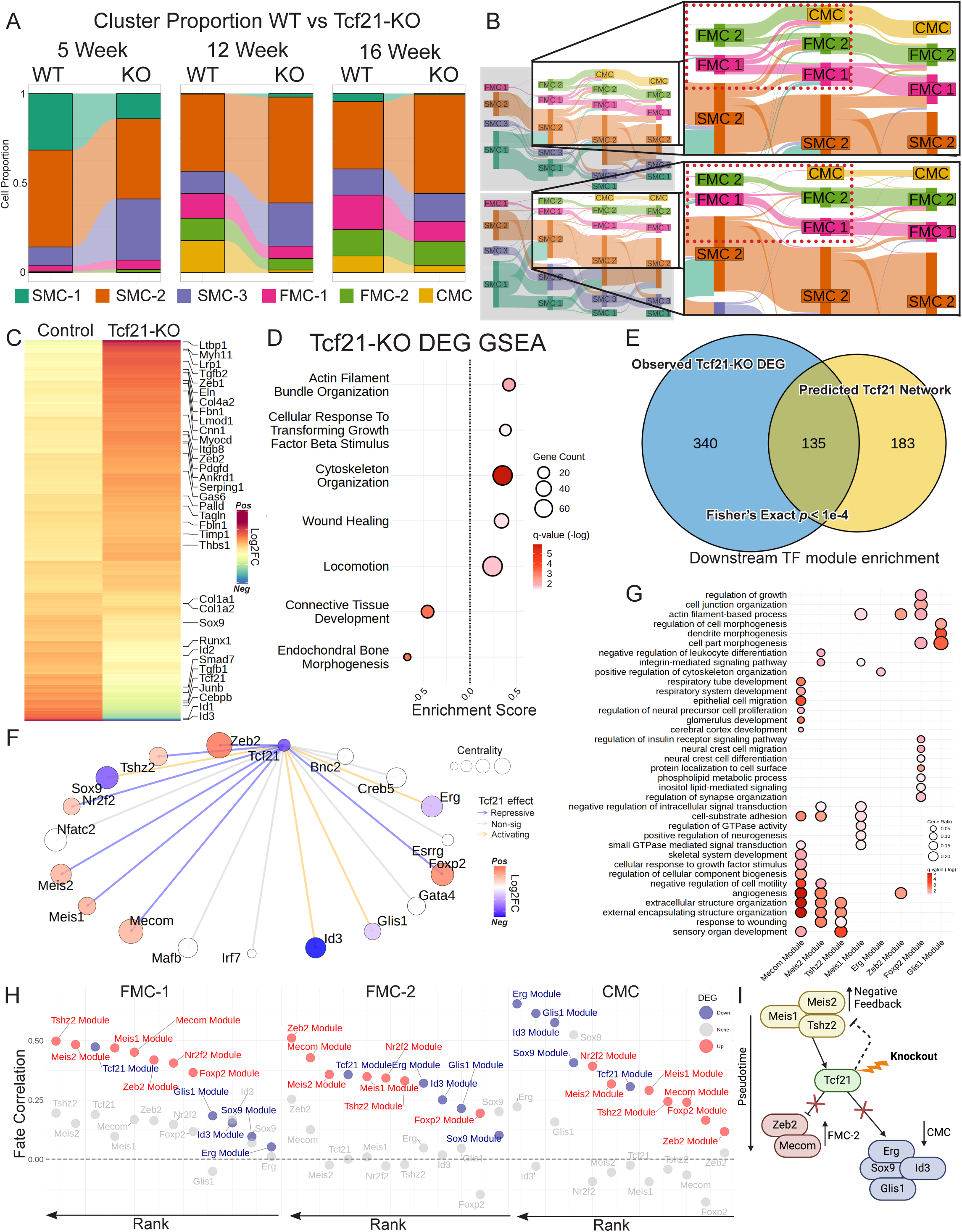
*Tcf21*-KO cells demonstrate differential cell fate trajectories and pathways associated with impaired phenotypic transitions. **A.** Comparison of cluster proportions in lineage traced SMC from control and *Tcf21*-KO mice matched by week of diet exposure. **B.** Sankey plot of control (top) and *Tcf21*-KO (bottom) WOT predicted transition probability from SMC to FMC and CMC populations. Band thickness represents relative transition probability from starting to end cell states. **C.** Heatmap of hierarchically clustered significantly differentially expressed genes between control and *Tcf21-*KO cells in the PseudoLate bins. **D.** GSEA (GO- biological process gene sets) of differentially expressed genes between control and *Tcf21*-KO cells in the PseudoLate bins. Size and color represent overlapping gene counts and enrichment q-value, respectively and x-axis shows GSEA enrichment scores. **E.** Over-representation analysis of observed *Tcf21*-KO DE genes overlap with the PANDO-predicted *Tcf21* TF network. **F.** Network showing predicted target TFs of *Tcf21*, observed *Tcf21*-KO effect on expression (edge, color), and network centrality of these target TFs (size). **G.** GO-BP enrichment of *Tcf21* target TF networks. **H.** Correlation for single cell WOT fate scores for FMC-1, FMC-2 and CMC fates with module scores for each *Tcf21* target TF network. Color represents TF expression directionality with *Tcf21*-KO. Control correlation of TF expression only with WOT fate scores in gray. **I.** Direct *Tcf21* network summarized in context of pseudotime.

WOT was used to investigate alterations in SMC trajectory transition probabilities in the context of *Tcf21*-KO **(Fig. 4B)**. Focusing on cell state changes in the SMC lineage traced cells, there was a notable decrease in the overall transition probabilities across cell states for *Tcf21*-KO. The SMC-2 contribution to both FMC-1 and FMC-2 was decreased in KO diseased tissues, as was the FMC-1 contribution to the FMC-2 phenotype, and there was a low level of transition for FMC-1 and FMC-2 to CMC **(Fig. 4B)**. Overall, there was evidence for a dramatic decrease in probability of transition to the CMC phenotype while transition to both the FMC-1 and FMC-2 phenotype was decreased but sustained. These decreased probabilities for SMC to FMC and FMC to CMC transitions in knockout mice accounted for the changes in SMC phenotype cell numbers but also highlighted the presence of compensatory processes which allowed continued phenotypic transition **(Fig. 4A).** As noted previously, at 16 weeks, the FMC-1 and FMC-2 states remained, suggesting that they represent a terminal phenotype rather than exclusively a transition to the CMC phenotype.

### *Tcf21*-KO DEGs demonstrate enrichment for timecourse predicted network and CAD genes

Analysis of differentially expressed genes (DEGs) in the *Tcf21*-KO compared to control mice using DESeq2 provided insight into associated gene programs. DEGs for the FMC-CMC transition for *Tcf21*-KO versus control, identified 965 genes **(S-Tables 9, 10)**. This list was enriched for TGFB family genes such as *Ltbp1* and *Tgfb2,* and numerous CAD GWAS genes including *Tgfb1, Zeb2*, *Lrp1, Palld, Col4a2, Lmod1*, and *Pdgfd* **(Fig. 4C)**. These DEGs were further analyzed with GSEA using GO-BP terms to gain insight into altered pathways and directionality of effect. We found “Cellular response to Tgfb stimulus” and “actin filament bundle organization” to have high positive DEG enrichment, consistent with *Tcf21* promoted effects and increased representation of “locomotion” and “wound healing” possibly representing compensatory processes given the overall decreased phenotypic transition phenotype **(Fig. 4D)**. Terms with a negative enrichment score (average downregulation with *Tcf21*-KO) were identified as suppressive for “connective tissue development” and “endochondral bone morphogenesis,” further suggesting that *Tcf21* target genes likely have a role promoting the CMC phenotype.

SMC lineage cell state changes and related cellular trajectories can be modeled through gene-gene interactions in a GRN. To build a *Tcf21* specific network we employed the regulatory network predicted from our PseudoLate control timecourse which spanned the majority of *Tcf21* activation in transitioning SMC. Moreover, network targets by this method are more likely to represent direct interactions given the conditional need for epigenetically accessible *Tcf21* binding sites to be linked to target genes. We compared *Tcf21* target genes identified with this network analysis with *Tcf21*-KO DEGs and found significant overlap of DE genes and network genes, with 135 of 318 network genes (42%) also showing differential expression with *Tcf21*-KO (Fishers exact test p<1e-4) **(Fig. 4E)**. Further, we applied GSEA using the predicted *Tcf21* network gene set ranked by TF-gene regulatory coefficient weights and observed a significant normalized enrichment score (NES = 1.24, p=0.031) of *Tcf21*-KO DEGs. These analyses showed congruence between distinct approaches and validated the utility of using predictive transcriptional modules from a comprehensive control dataset to infer perturbed pathways.

We leveraged the predictive pathway ability of inferred networks to augment the functional characterization of the *Tcf21* signaling network. We selected differentially expressed TFs within the *Tcf21* GRN and integrated these TF-centered GRNs to create a validated *Tcf21*-TF sub-network and find both repressed and activated TFs that are regulated by *Tcf21* **(Fig. 4F)**. We identified downstream TF networks with differential KO module scores and performed functional enrichment on this subset of TF networks to visualize the pathways affected. We found multiple TFs including *Foxp2*, *Mecom*, *Zeb2*, *Meis2*, *Mecom* and *Tshz2* predicted to be involved in cell migration, angiogenesis and extracellular organization processes, while genes such as *Meis1*, *Zeb2* and *Foxp2* were also involved in contractile processes **(Fig. 4G).**

We computed TF fate correlations by comparing WOT transition probability with cell-level TF module scores and using TF-only expression as a control comparison **(Fig. 4H)**. Interestingly, top modules that exhibit high correlation with FMC fates also have their central TF upregulated with *Tcf21*-KO, suggesting compensatory roles alongside *Tcf21* given our observation of overall decreased FMC proportions. For example *Zeb2*, previously shown to drive phenotypic transition ^13^, was increased in *Tcf21*-KO, and its later average TF- pseudotime suggested that its role is downstream of *Tcf21* towards the FMC-2 fate. In contrast, TFs such as *Meis1/2* have earlier average TF-pseudotime, suggesting compensatory upregulation possibly via feedback mechanisms. Conversely, top TF modules correlated with the CMC fate, including the known ossification regulator *Sox9*, showed decreased expression upon *Tcf21*-KO **(Fig. 4I)**.

### Integrative epigenetic analysis of Tcf21-KO identifies TCF21-TEAD epigenetic interactions modifying CAD GWAS genes

Previous studies have shown that *Tcf21* interacts with histone deacetylases to broadly alter the *in vitro* epigenetic landscape of human coronary artery SMC (HCASMC) ^51^ but its effects *in vivo* have not been explored. We observed widespread motif deviation changes upon *Tcf21*-KO when visualizing differential ChromVar scores across pseudotime and generated differential ChromVar scores on a by cluster basis **(Fig. 5A, S-Tables 11,12)**. We further used Jensen-Shannon divergence (JSD) scores to identify differentially deviated TF motifs across pseudotime between control and *Tcf21*-KO ChromVar distributions, finding significant differences in many core TF motifs including Zeb, Klf, Runx1/2, AP-1, Srf, and Ctcf, while Tcf21 was borderline significant (padj = 0.10) **(Fig. 5B).** *TCF21* HCASMC ChIPseq was reprocessed from Zhao et al.^52^ and showed significant enrichment in TEAD and CEBP TF binding motifs within TCF21 peaks (**Fig 5C, S-Fig. 9A**) in addition to TCF21 and AP-1^51^. When these ChromVar scores were visualized along pseudotime in the mouse time course, there was enhancement of Tead1 motif accessibility with loss of *Tcf21* while *Cebpb* shared a pattern similar to *Tcf21*, showing decreased accessibility with *Tcf21*-KO (**S-Fig. 9B**). We performed additional ChIPseq in HCASMC for *TEAD1* and confirmed significant overlap with *TCF21* **(Fig. 5C**, Fisher’s exact test p<1e-4). Pathway analysis of shared peaks using GREAT predicted shared biological functions related to inflammation, apoptosis, TNF, and TGFB **(Fig. 5D)**. Peaks shared by *TCF21* and *CEBPB* were enriched for “cell adhesion”, “ERK signaling”, and “cell motility terms” **(S-Fig. 9C, D**, Fishers exact test p<1e-4). Genome wide colocalization of *TCF21* and *TEAD1* motifs was identified by comparing the location of *TEAD1* motifs in *TCF21* ChIPseq peaks and *TCF21* motifs in *TEAD1* peaks **(S-Fig. 9E).** Interestingly, enrichment analysis for GWAS SNPs at *TCF21*-*TEAD1* co-bound loci using the GWASAnalytics package revealed a greater odds ratio for CAD (**Fig. 5E**) compared to both TF alone and *TCF21* plus *CEBPB* **(S-Fig. 9F-I)**. This *TCF21*-*TEAD1* relationship corresponded with murine trajectory analysis nominating *Tead1* as an FMC-2 driver **(Fig. 2L)** and making it an intriguing *Tcf21*-interacting partner for further study.

**Figure 5.**
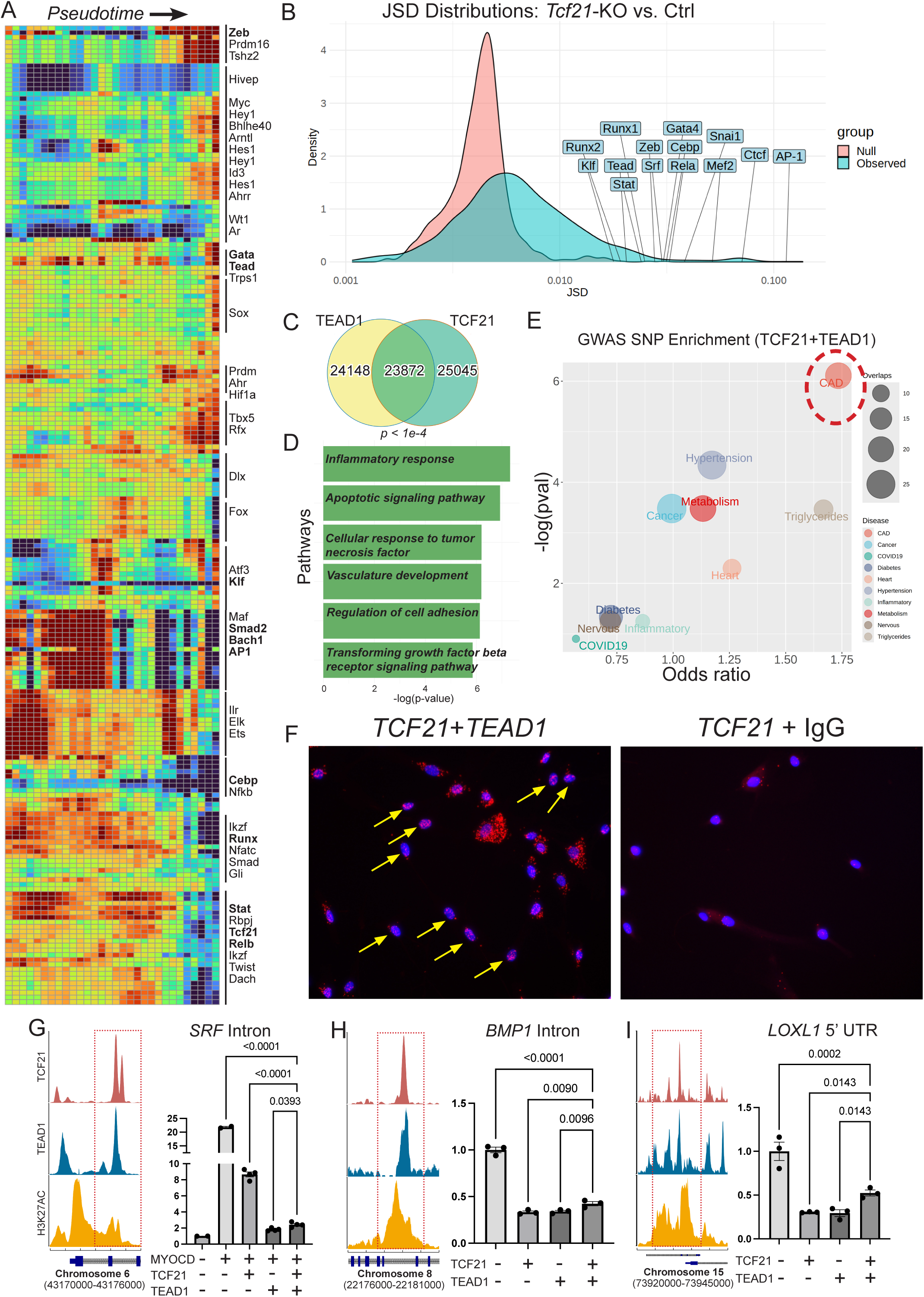
*TCF21* mediates genome wide epigenetic effects and co-localizes with *TEAD1* to epigenetically regulate SMC genes involved in CAD risk and guide cell state transitions. **A.** Hierarchically clustered heatmap of differential ChromVar scores (*Tcf21*-KO vs. control) labeled with selected representative TF motifs **B.** Jensen-Shannon divergence of observed *Tcf21*-KO vs. control ChromVar scores showing motifs demonstrating significant probability divergence between conditions. Control JSD is calculated by randomly splitting control dataset into two equal distributions and calculating the JSD between these distributions. **C.** ChIPseq peak overlap between TCF21 and TEAD1. **D.** Representative pathways identified from shared ChIPseq peaks using Genomic Regions Enrichment of Annotations Tool (GREAT). **E.** GWASAnalytics analysis of GWAS SNP (EMBL-EBI GWAS Catalog) enrichment of overlapping ChIPseq peaks between *TCF21* and *TEAD1* showing highest enrichment for CAD traits. **F.** Proximity ligation assay showing nuclear fluorescent signal enrichment in the *TCF21*+*TEAD1* antibody group compared with TCF21+IgG control. **G, H, I.** Dual luciferase assay for overlapping regulatory regions (*SRF* Intron, *BMP1* Intron, *LOXL1* Promoter) for *TCF21* and *TEAD1* demonstrating competitive repression and epigenetic fine-tuning of enhancer/promoter activity between regulatory elements.

We further examined the shared genomic patterns between *TCF21* and *TEAD1* by partitioning their shared binding loci into *TCF21*+*TEAD1* or *TCF21* only loci. We observed greater enhancer profiles for *TCF21*+*TEAD1* shared binding sites compared to *TCF21* only, as indicated by overlap with H3K27ac **(S-Fig. 9L).** *TCF21*+*TEAD1* shared binding sites were located further from the TSS regions (**S-Fig. 9M)**, consistent with enhancer co-localization. Further, pathway analysis of putative genes identified in *TEAD1*/*TCF21*/*H3K27* regions by GREAT revealed enriched pathway keywords including “differentiation, “development” and “endopeptidase regulation” while *TCF21*-only+H3K27 region genes showed enrichment for “immune,” “viral” and “neutrophil” related keywords (**S-Fig. 9N, O**).

We further investigated physical and functional interaction of these two TFs. Proximity ligation assays found that *TCF21* and *TEAD1* co-localized in the nucleus, suggesting direct protein-protein interaction (**Fig. 5F**). To examine the functional interactions between *TCF21* and *TEAD1*, we performed dual luciferase reporter gene transfection assays with A7r5 rat smooth muscle cells on a shared enhancer residing in an intron of the SRF gene, a master regulator of lineage contractile gene expression^36^, and two additional enhancers in CAD loci encoding ECM effectors of TGFB signaling, *BMP1* and *LOXL1*. For the *SRF* enhancer, we showed that normal activation by *SRF* binding partner *MYOCD* was highly suppressed by *TCF21* and to a greater degree by *TEAD1* alone (**Fig. 5G**). There was an intermediate reporter activity when *TCF21* and *TEAD1* were both transfected, suggesting a competitive interaction between these TFs. Also, for the *BMP1* and *LOXL1* enhancers, both TFs showed repressor activity, but again, intermediate suppression when both were expressed in the same cells, suggesting competition (**Fig. 5H, I**). Taken together these data suggest that *TCF21* and *TEAD1* directly interact at shared regions across the genome to epigenetically regulate transcription.

### Genetic CAD risk attributable to the SMC lineage is mediated by *TCF21* through early phenotypic transition network rewiring

Single cell methods have previously nominated genetic risk signals to have a unique high enrichment in SMC^8^. We investigated the relative disease related significance of our SMC cell states, using the scDRS algorithm^5,53^. At the single cell level, we leveraged scDRS to integrate gene expression and GWAS gene z-score weights from MAGMA to generate disease relevance scores for each cell type. We also computed GWAS risk-weighted average expression using scDRS gene weights multiplied by its average expression and inferred directionality using updated heritability adjusted S-PrediXcan modeling^54^ **(S-Table 13).** We identified greater averaged expression of CAD risk genes in FMC-2 and CMC relative to FMC-1 and SMC. Moreover, scDRS found FMC-2 to be significantly enriched for CAD risk genes **(Fig. 6A)** Interestingly, neither the transcriptionally similar FMC-1 nor the calcification related CMC showed significant evidence for disease risk association by scDRS, prompting further investigation of the distinguishing expression patterns of these transition phenotypes.

**Figure 6.**
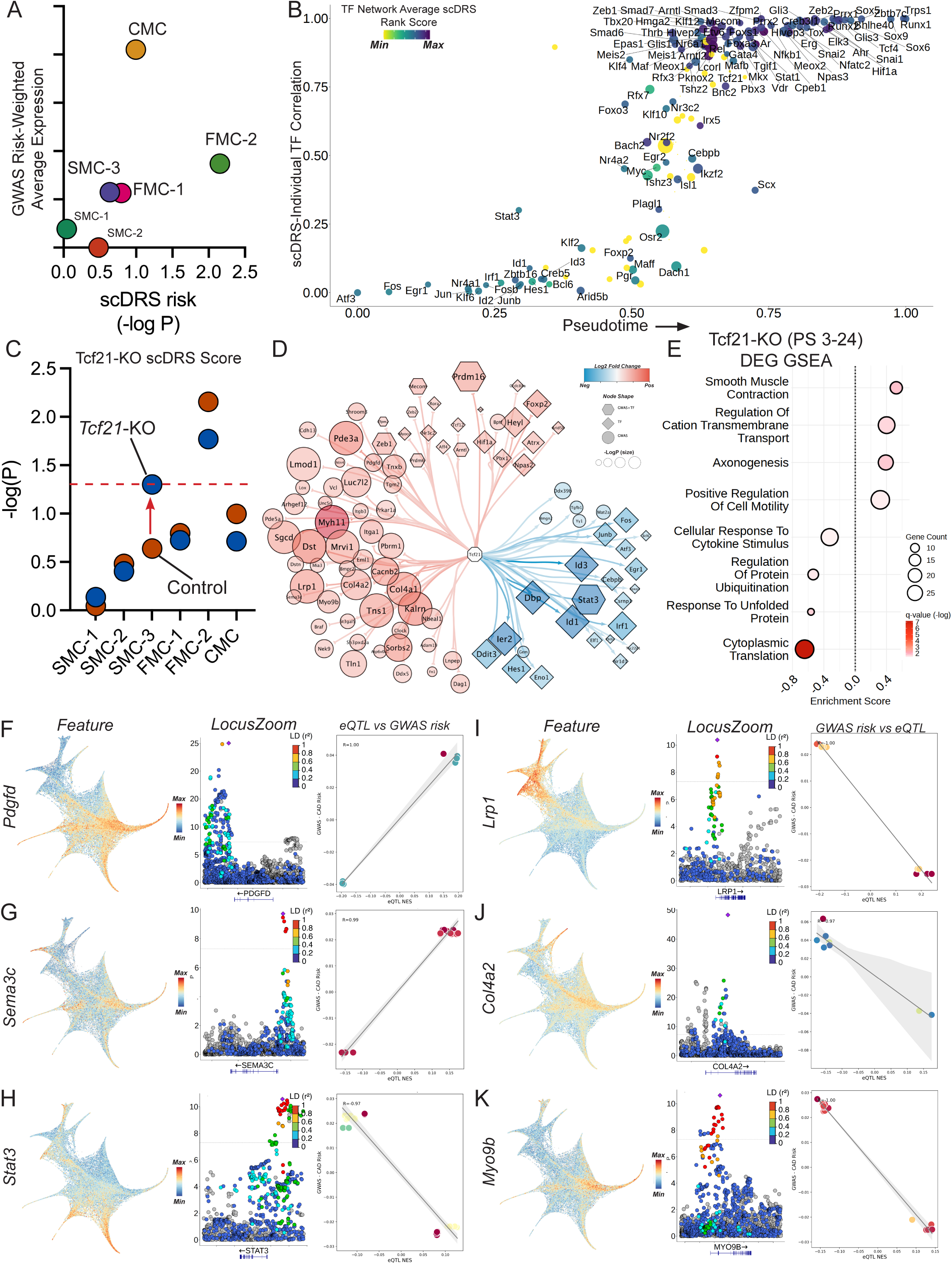
CAD GWAS integration with scRNASeq data identifies baseline disease risk in SMC transition cell states and and changes in disease gene expressionwith *Tcf21*-KO. **A.** Scatter plot of CAD GWAS risk-weighted average expression of disease genes based on MAGMA gene weights and PrediXcan estimated directionality versus GWAS gene enrichment significance calculated by scDRS for each SMC cell state. **B.** Scatter plot of individual TF and scDRS score expression correlation across pseudotime. Color represents averaged scDRS gene rank for TF gene modules and size represents overlap with GWAS genes. **C.** Change in scDRS significance per cluster with *Tcf21*-KO, dotted line indicating nominal significance (p = 0.05). **D.** Summarized *Tcf21* PseudoEarly DEG network filtered for putative GWAS genes and TFs. Color scale and node shape indicate differential expression and gene type, respectively. **E.** GSEA(GO-biological process gene sets) of differentially expressed genes between *Tcf21*-KO and control SMC in the PseudoEarly bins. Size and color represent overlapping gene counts and enrichment q-value, respectively. The x-axis shows GSEA enrichment scores. **F-K.** Gene expression feature plots and correlation of arterial tissue eQTL NES to GWAS odds risk for overlapping single nucleotide polymorphisms of selected *Tcf21*-KO differentially expressed genes from PseudoEarly bins with shared **(F-H)** or opposing **(I-K)** expression-disease risk directionality with *Tcf21*.

We used the TF networks identified from the control timecourse (Methods) to ask how individual TFs and their networks associate with CAD risk. First, we synthesized full TF network modules by assimilating all unique target connections from PseudoEarly and PseudoLate predicted regulatory networks for available TFs. Second, we extracted gene-level scDRS weights to derive normalized TF-scDRS correlations. Third, we averaged gene-scDRS correlation ranks of all genes within each TF network module to create a TF network average scDRS rank. At the network level, we observed TFs with higher average scDRS rank in the SMC to FMC pseudotime such as *Tcf21*, *Nkfb1*, *Runx, Zeb, Hif* and *Smad* factors **(Fig. 6B, S-Table 14)**. Many of these core TFs also overlap with *in silico* TF perturbation predictions **(Fig. 3F-G)**. Moreover, this method nominated novel TFs with network enrichment of CAD risk genes that were postulated to drive the SMC-FMC transition, including *Arntl*, *Prrx1*, *Tshz2*, and *Mecom* **(S-Table 8***) or* the FMC-CMC transition such as *Trps1*, *Zbtb7c*, *Snai1*, and *Sox5/6/9*.

To further dissect the regulatory relationships of CAD GWAS genes in phenotypic transition, we examined the scDRS enrichment in *Tcf21*-KO and found an increase in SMC-3 scDRS z-score, meeting nominal significance **(Fig. 6C)**. This shift in scDRS score implicated the ability of *Tcf21* to coordinate CAD GWAS genes early in the phenotypic transition timeline. In addition, this observation was in agreement with *Tcf21* expression showing SMC-3 enrichment (**Fig. 1C**) and demonstrating a basal level of accessible *Tcf21* TF binding sites in early pseudotime (**Fig. 3B**). We then examined the DEGs in PseudoEarly bins and observed significant overlap with a curated set of putative GWAS genes (63/640, *p* < 0.0001) **(Fig. 6D)**. GSEA enrichment of DEGs not only revealed similar increased contractile processes with *Tcf21*-KO but also highlighted a decreased response to stress signals such as ‘cytokine stimulus’ and ‘unfolded protein response’ **(Fig. 6E)**. Integrating these genes with human genetic signals, we focused on the differential expression of genes that shared GWAS disease risk-eQTL correlation with *Tcf21*. For example, increases in proliferative factors such as *PDGFD* and *SEMA3C* or decreases in inflammatory transcription factor *STAT3* were associated with increased CAD risk **(Fig 6. F-H)**. Conversely, there was also an enrichment of genes that were correlated with decreased CAD risk such as *LRP1* which has multifaceted coronary disease implications, *COL4A2* which promotes basement membrane integrity or *MYO9B* that modulates vascular wound repair **(Fig 6. I-K)**. Together, these relationships suggest a broad role for *TCF21* in promoting risk through coordinated regulation of CAD GWAS genes in the phenotypic transition of disease SMC.

## Discussion

We have conducted a comprehensive single cell study to investigate the molecular trajectory of SMC phenotypic transitions during atherosclerosis using a combination of multi-modal single cell sequencing at multiple timepoints, key marker in situ hybridization, and trajectory modeling. These data provide transcriptomic, epigenomic and cellular lesion anatomical data characterizing two different FMC populations. FMC-1 arise first by 5 weeks of diet exposure, express inflammatory and immune markers while FMC-2 accumulation accelerates weeks later and is characterized by extracellular matrix, lipid handling, osteoblast progenitor expression profiles and a greater correlation with contractile marker expression compared to FMC-1. Trajectory modeling suggested FMC-1 contribute to FMC-2, but both phenotypes arise primarily from a modulating group of cells that maintain classical SMC contractile marker expression. FMC-1 are localized to the media and to the fibrous cap, suggesting their involvement in migration, while FMC-2 are identified primarily at the fibrous cap and intimal plaque. Both FMC contribute to CMC transition cells and are likely the sole source of these endochondral bone-like phenotype cells.

Our comprehensive multi-omic dataset has enabled the profiling of TF motif accessibility gradients across time and leveraged these data to generate regulatory networks, prioritize key driver genes and evaluate their functional pathways. Analysis using both WOT and pseudotime suggested that both FMC-1 and FMC-2 phenotypes transition probabilities stabilize in mature lesions and are thus more likely to represent terminal phenotypes rather than a transition state. This possibility needs to be further investigated with longer timepoints for diet exposure, but it is an attractive possibility, following the hypothesis that protective cells transition primarily to stable FMC and disease promoting cells undergo further transition to the CMC lineage.

We further integrated trajectory analysis and in silico TF perturbation to identify critical factors which establish cell state identity through their ability to physically access genomic regulatory regions. For example, our aggregated SMC to FMC transition analysis **(Fig. 3H, I)** showed extensive overlap of top factors known to affect SMC phenotypic transition to FMC or CMC such as *Tcf21*, *Ar*^55^, *Runx1/2*^49,56^, *Zeb2*^13^, *Smad3*^12^, and *Klf4*^11^ while also nominating novel TFs such as *Arntl, Mecom*, *Prrx1*, *Trps1, Zbtb7c,* and a variety of HIF-related factors that participate in divergent functional roles for future study.

Among the prioritized TFs, *Tcf21* emerged as a compelling candidate given its top rank in predicted effects on phenotypic transition as well as our previous work identifying it as a causal CAD GWAS gene and providing human genetic evidence for its CAD risk inhibition^16^. First generation scRNAseq studies have shown that *Tcf21* loss was associated with decreased fibroblast-like SMC lineage cells that we termed fibromyocytes and histology showed decreased SMC migration from the media and decreased contribution to the fibrous cap. Therefore, our focused timecourse single cell study in the *Tcf21*-KO mouse model with enhanced scRNAseq chemistry, greater transcriptomic depth, and linked chromatin accessibility data allowed further analysis of cellular trajectories and phenotypic transitions altered by *Tcf21* loss. These analyses identified novel TF networks directly altered with *Tcf21*-KO as well as interacting epigenetic factors such as *Tead1* which together with *Tcf21*, antagonize the differentiated SMC cell-fate and fine-tune cellular TGFB response.

We also found *Tcf21* to be enriched in SMC-3 cells that emanate in part from the SHF, and the *Tcf21*- KO mouse showed a dramatic 3-fold expansion in the *Tnnt2* expressing SMC-3 cells after only 5 weeks of diet. This observation suggests that *Tcf21* suppresses transition, and possibly migration, of cells from this region.

SMC derived from the SHF give rise to the proximal aortic wall and to the adjacent outflow tract and exhibit well-recognized embryonic lineage specific responses to critical signaling pathways^44^ such as TGFB^57^, PDGFD^58^, and NFkB^59^. Further, increased *Tcf21* expression was identified after disease initiation in the FMC-1 where its expression was noted to be inversely correlated with expression of *Acta2* and other contractile markers, consistent with our previous findings that *Tcf21* suppresses SMC lineage marker genes through direct transcriptional mechanisms that block *MYOCD-SRF* mediated transcription of lineage markers^36^. We made the new observation that loss of *Tcf21* resulted in decreased transition to the CMC phenotype. This was demonstrated by decreased lineage traced CMC proportions, reduced WOT calculated transition probabilities and also supported by the relative importance assigned to *Tcf21* as an enriched TF in WOT analysis of cells fated to become CMC **(Fig. 2F)**. Given that human genetic data is consistent with *TCF21* having a protective role toward CAD risk, these new findings suggest that the most relevant mechanism by which SMC relevant CAD genes affect human disease risk may be primarily related to the ability of SMC lineage cells to contribute to the FMC cellular phenotypes. In this case, the promotion of CMC number cells would be secondary to the increased FMC transition to CMC, akin to a mass action effect that does not negatively impact disease risk.

We and others have used CAD GWAS findings along with gene expression or chromatin accessibility data to show that much of the risk for CAD resides in the coronary vascular SMC lineage^5,8,9,53^. Our high-resolution dataset extends upon previous observations and identifies FMC-2 cells as the cell state harboring the greatest CAD risk based on disease gene expression integration with scDRS and PrediXcan and suggests that the FMC-2 cell state is a determining mechanism of human CAD risk **(Fig. 6A)**. In situ RNA staining localizes the FMC-2 population to both the fibrous cap (*Notch3, Thbs1*) and the intimal plaque (*Fbln2, Ttc9)*.

Further molecular analysis finds it at the juncture of critical gene modules for senescence and apoptosis while expressing numerous CAD associated genes that we have linked to atherosclerosis, including *ZEB2* and *SMAD3*^12,13^. In addition, examination of how *Tcf21*-KO alters CAD risk associated genes finds the largest shift in SMC-3, again highlighting this unique cell state with SHF markers. Using this ‘differential’ scDRS score leverages the power of GWAS genes to suggest that although expressed at lower levels, the cell state determining effects of *Tcf21* likely occur early and are critical to prime cells towards phenotypic transition.

We have created these high order genomic data sets to map the epigenetic and transcriptomic mechanisms that mediate the SMC lineage transitions and their contribution to disease risk. This study focused on genes that are expressed by transition SMC and linked to phenotypic cell state changes through TFs and other high content signaling molecular pathways that mediate disease trajectories. We have identified a number of such genes that reside in CAD associated loci and are candidates for causal relationships with SMC transitions **(S-Table 8).** These genes were not validated as causal with genome editing or animal model studies, as such efforts are beyond the scope of the present work, but many show colocalization of expression quantitative trait and CAD associated variation suggesting causality. These example candidate genes allow a number of observations relating SMC transition and CAD gene association. Although appreciated previously, it is clear that numerous TGFB pathway genes represent a significant component of disease risk in relation to SMC phenotypic transitions. Other represented gene programs include chondrogenesis, hypoxia response, vascular development, and epithelial mesenchymal transition, in addition to proliferation and migration.

Importantly, while most processes have expression across multiple SMC lineage phenotypes, they are all enriched in FMC-2 cells. Finally, although it will require extensive additional mechanistic experimentation, these data suggest that both disease promoting as well as disease inhibiting factors are associated with the cell state changes that SMC undergo in the context of disease stresses.

## Supplemental Figure Legends

**Supp Fig 1.**
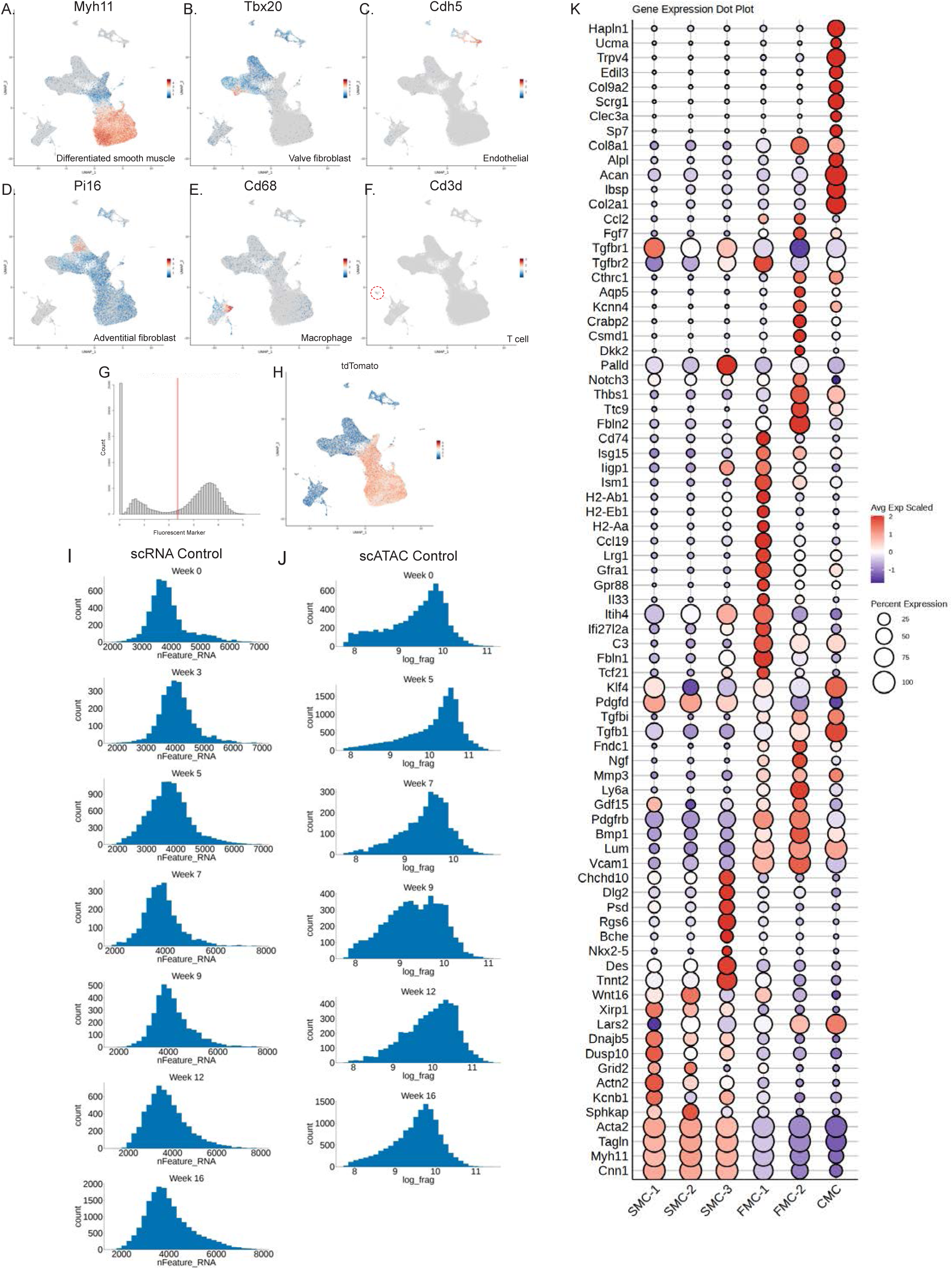
**A-F.** Gene expression feature plot for key features defining cell states for global (tdTomato and non-tdTomato) single cell capture. **G.** Histogram of expression counts for tdTomato and red-line indicating cutoff determined by logistic regression for tdTomato positive cells. **H**. Gene expression feature plot for tdTomato. **I.** Single cell RNA quality control metrics for control cells **J**. Single cell ATAC quality control metrics for control cells. **K.** Dotplot showing marker gene expression across SMC cell states. Color and size represent average scaled expression and percentage of cells expressing marker gene within cluster, respectively.

**Supp Fig 2.**
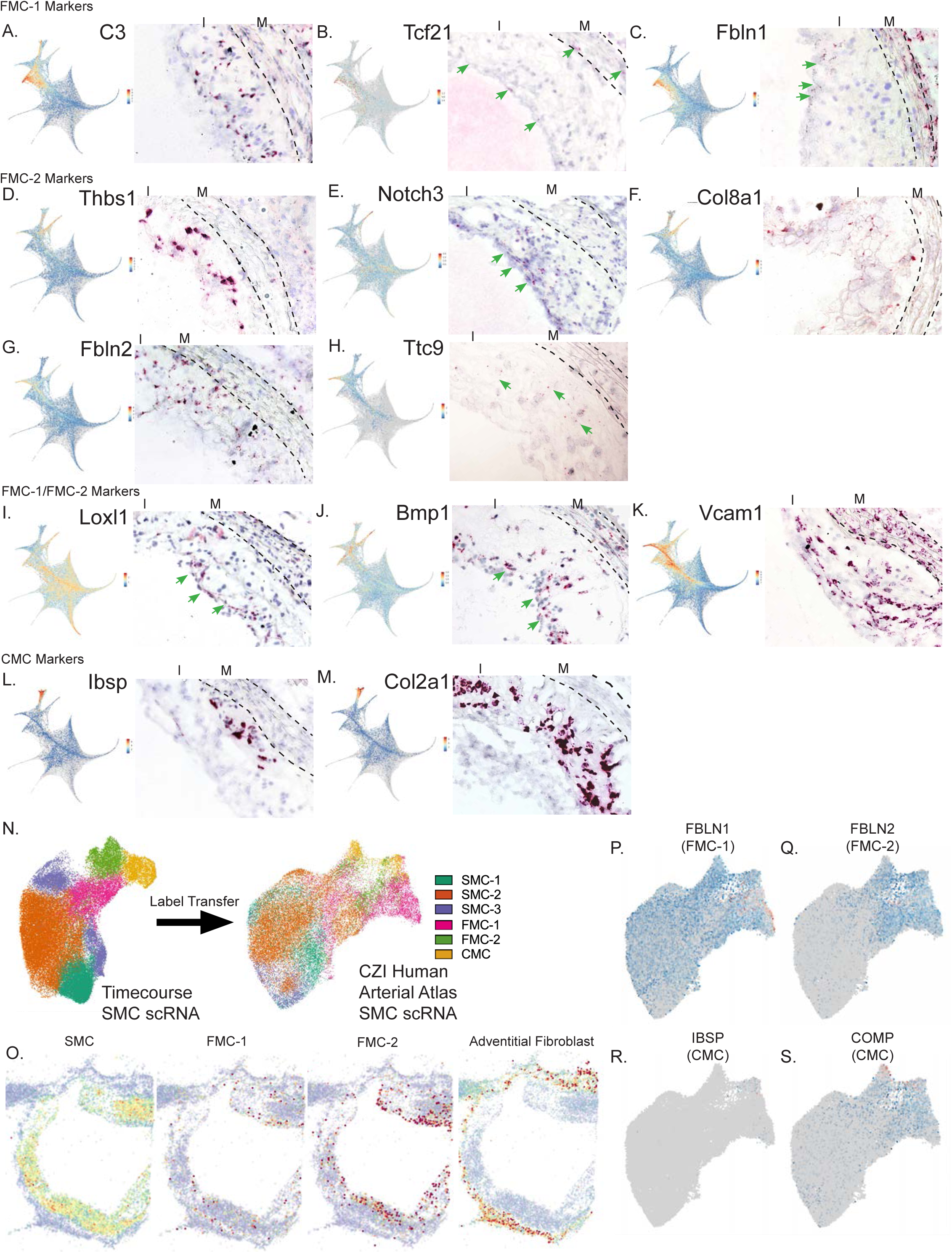
**A-C** Gene expression feature plot (left) and RNAScope staining (right) for FMC-1 markers. **D-H.** Gene expression feature plot (left) and RNAScope staining (right) for FMC-2 markers. **I-K.** Gene expression feature plot (left) and RNAScope staining (right) for pan-FMC markers. **L-M.** Gene expression feature plot (left) and RNAScope staining (right) of CMC markers. **N.** Murine SMC cell label transfer to CZI human arterial atlas SMC. **O.** Human coronary artery Slide-Seq showing spatial localization of cell state features based on predicted labels from CZI human arterial atlas SMC scRNA. **P-S.** Gene expression feature plot for representative cell state marker genes.

**Supp Fig 3.**
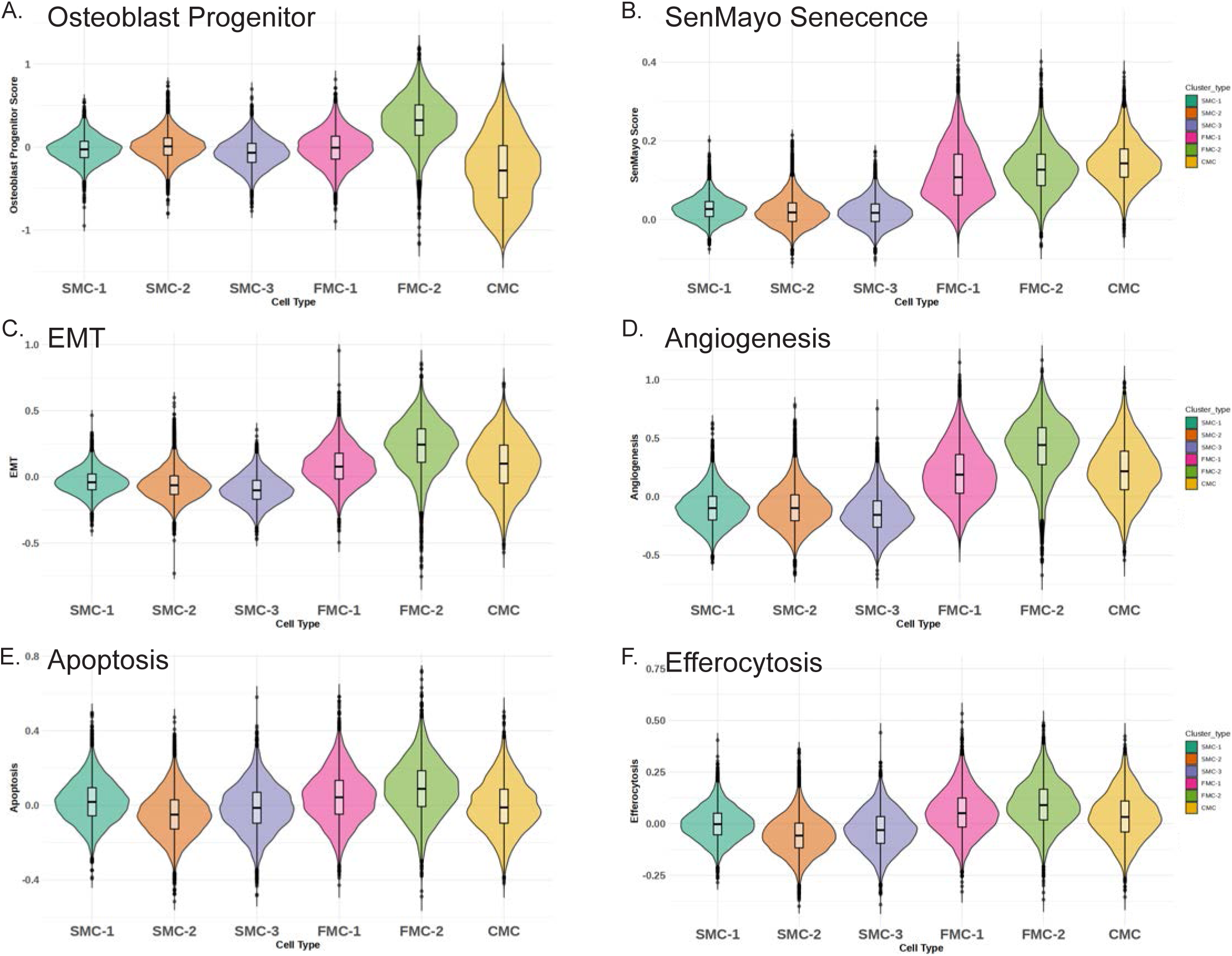
**A-F.** Gene module scores for osteoblast progenitor score, SenMayo Senescence score, EMT (epithelial mesenchymal transition), angiogenesis,, apoptosis, and efferocytosis split by cell state.

**Supp Fig 4.**
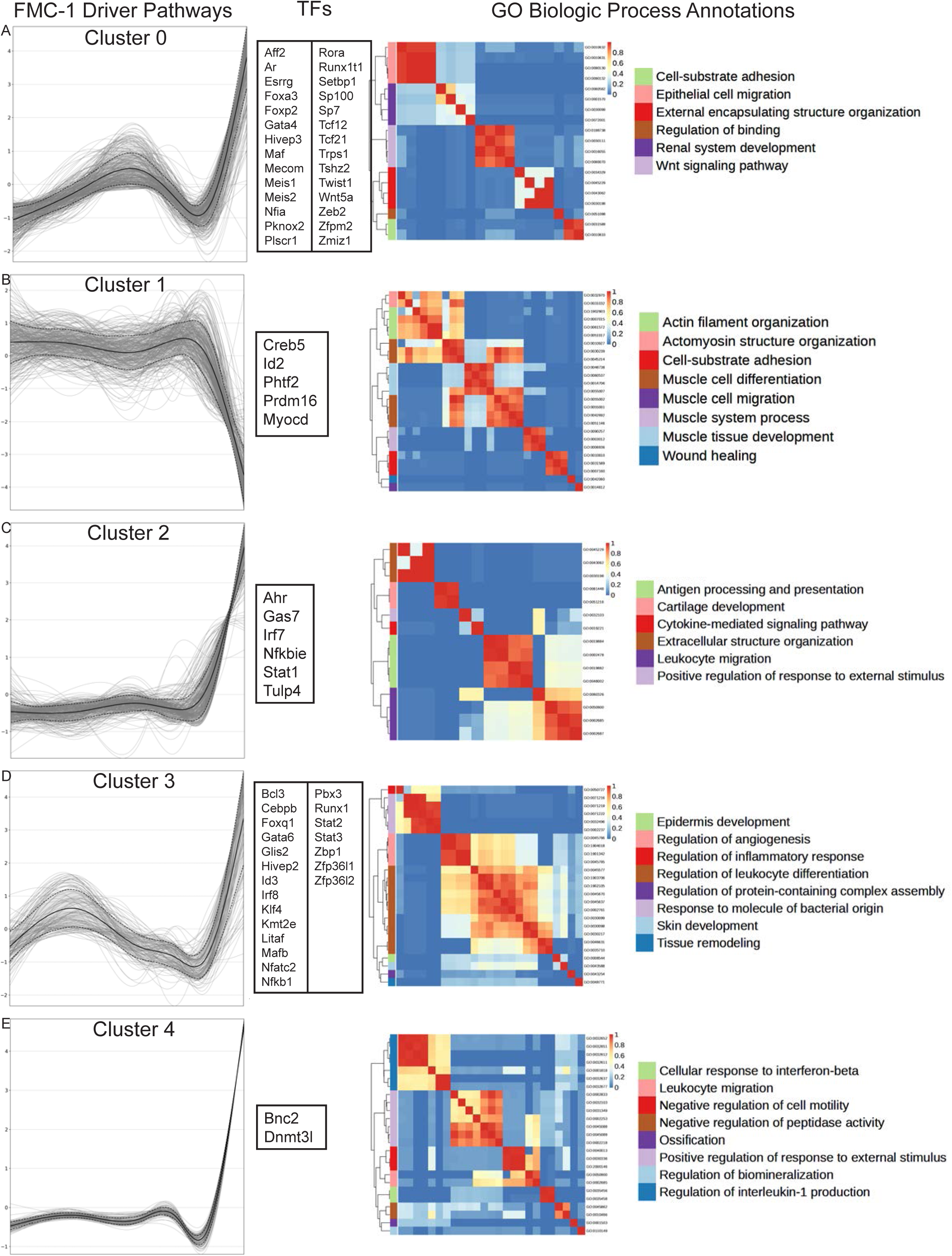
**A-E.** CellRank2 summarized expression trends for FMC-1 showing clustered gene expression pattern across pseudotime (right), TFs within this gene cluster (middle), and GO enrichment of cluster genes (left).

**Supp Fig 5.**
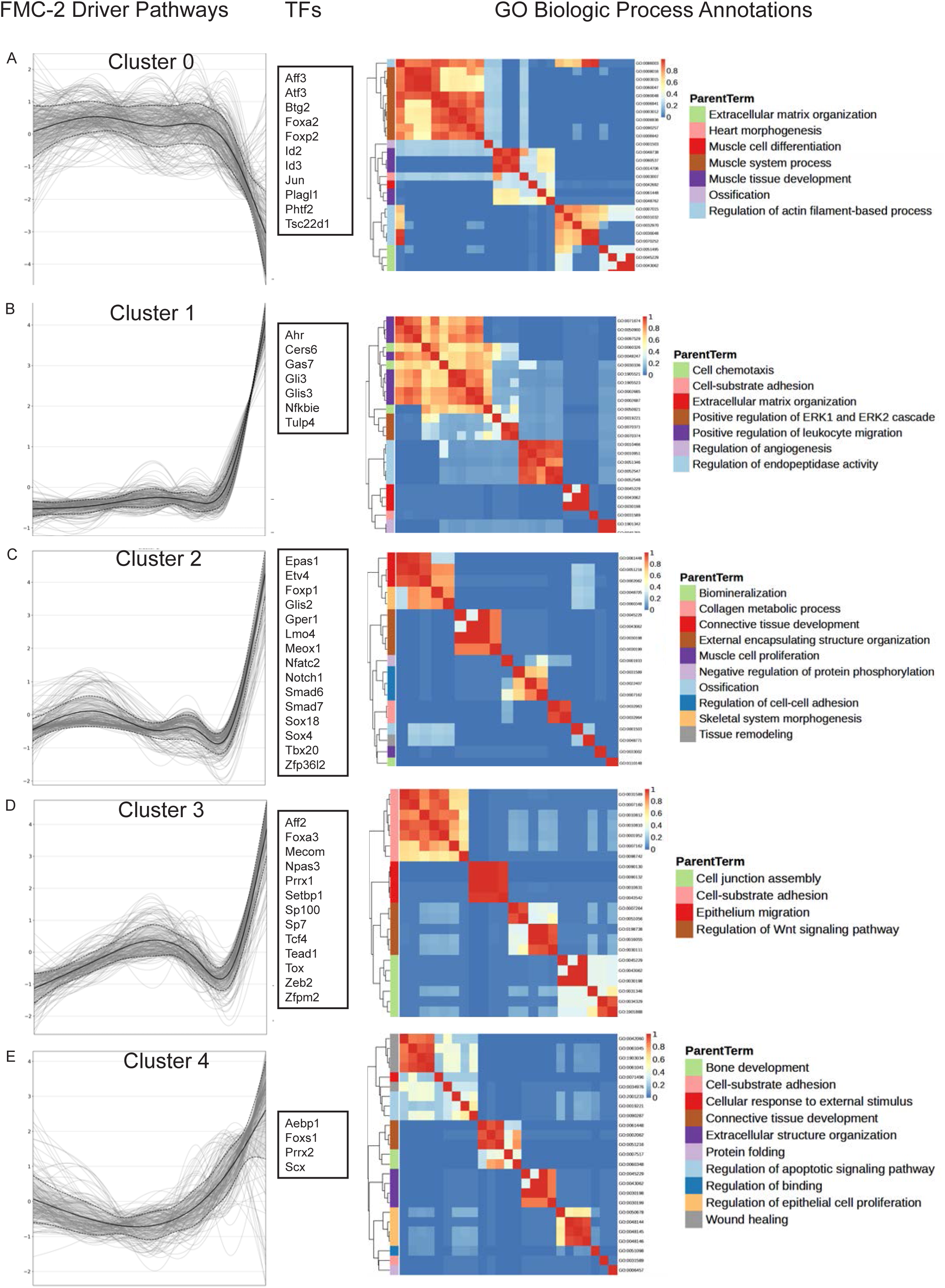
**A-E.** CellRank2 summarized expression trends for FMC-2 showing clustered gene expression pattern across pseudotime (right), TFs within this gene cluster (middle), and GO enrichment of cluster genes (left).

**Supp Fig 6.**
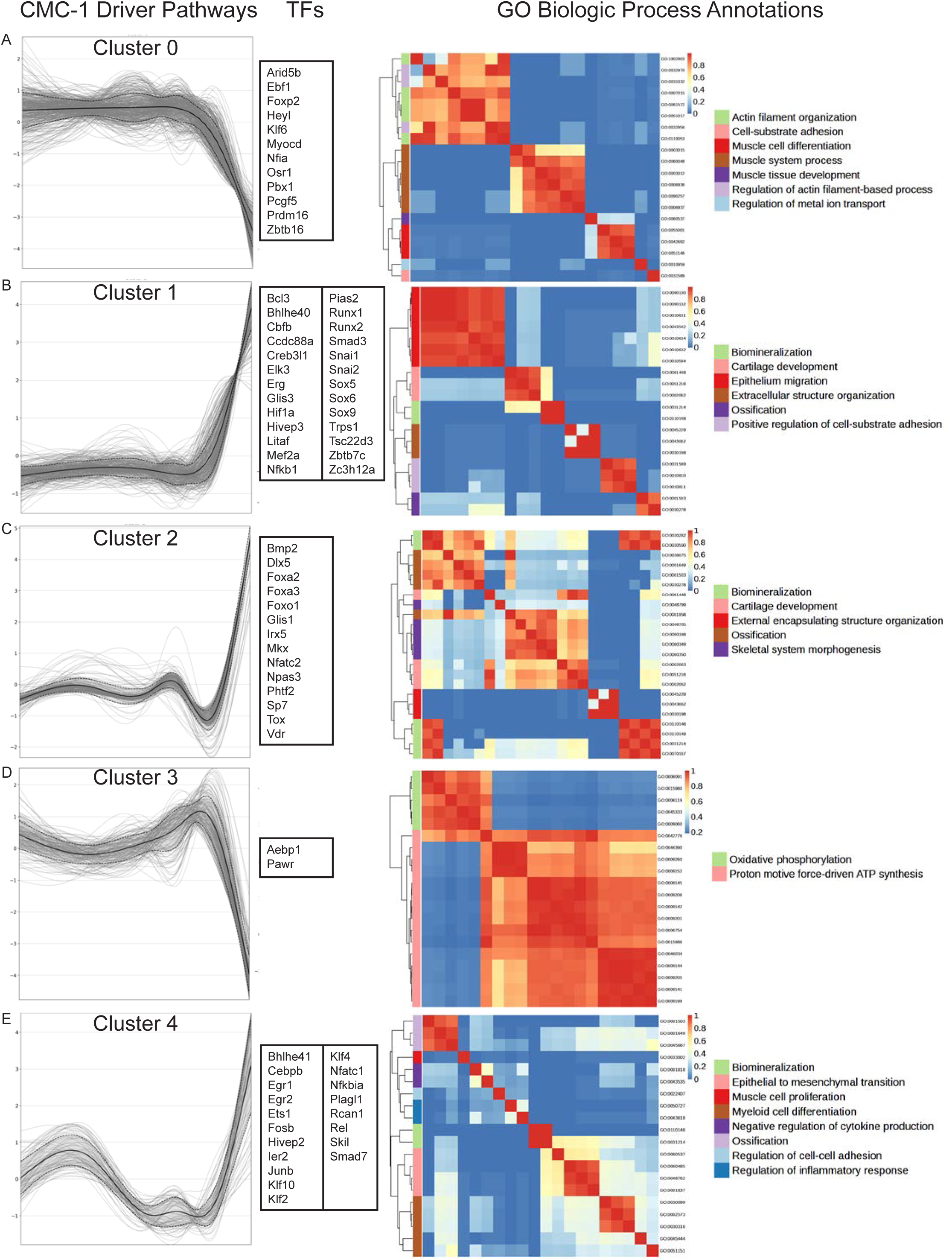
**A-E.** CellRank2 summarized expression trends for CMC-1 showing clustered gene expression pattern across pseudotime (right), TFs within this gene cluster (middle), and GO enrichment of cluster genes (left).

**Supp Fig 7.**
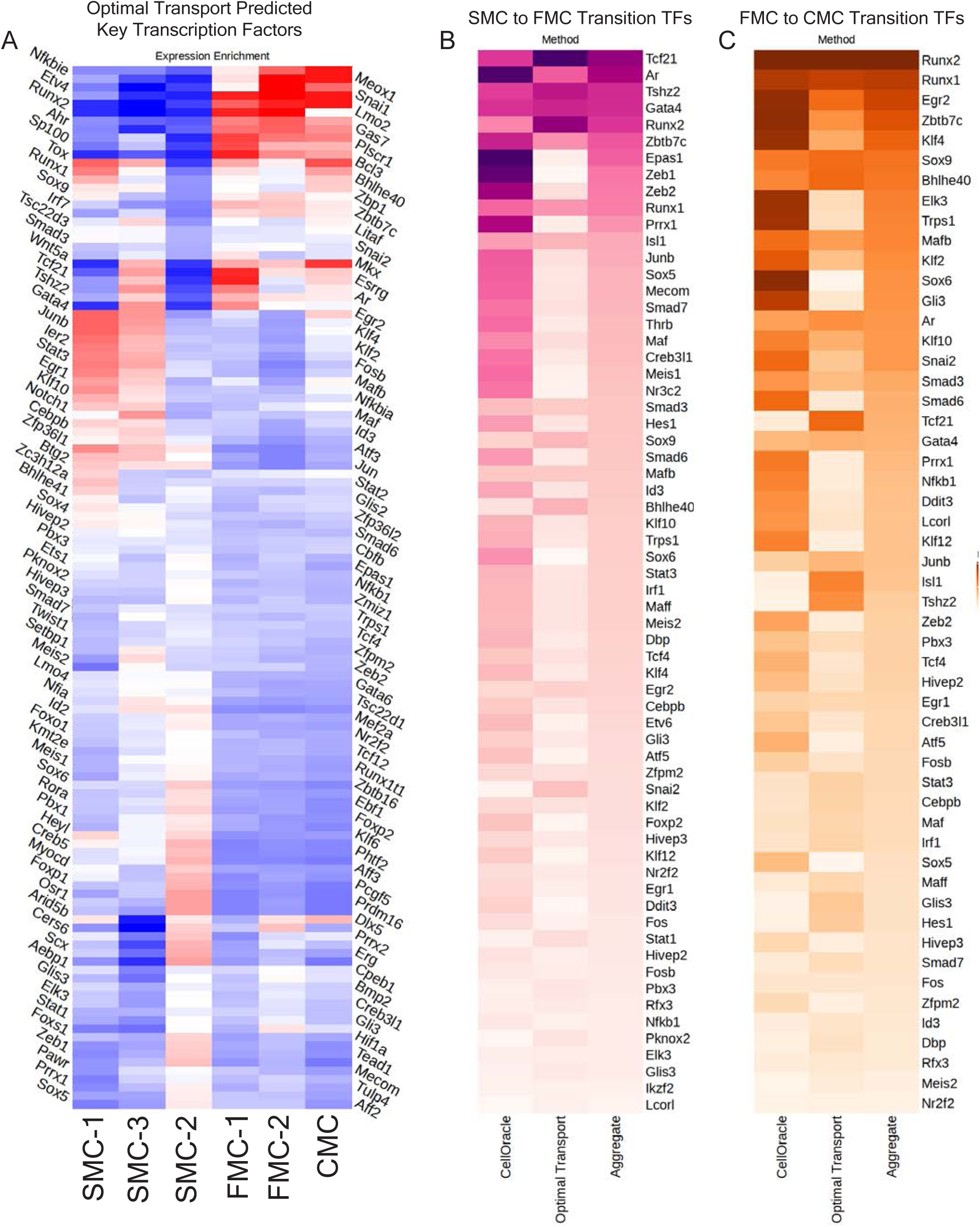
**A.** Unfiltered (including TFs without over-represented ATAC motifs) transcription factor enrichment for cells fated to become each cluster-type calculated by WOT based on week 5 to week 12 transition. **B-C.** Unfiltered (including TFs without over-represented ATAC motifs) ranking of top TFs predicted to affect the SMC to FMC or FMC to CMC transition using 1) CellOracle in silico perturbation scores, 2) Optimal Transport derived fated cell TF enrichment, or 3) aggregated ranking across both methods.

**Supp Fig 8.**
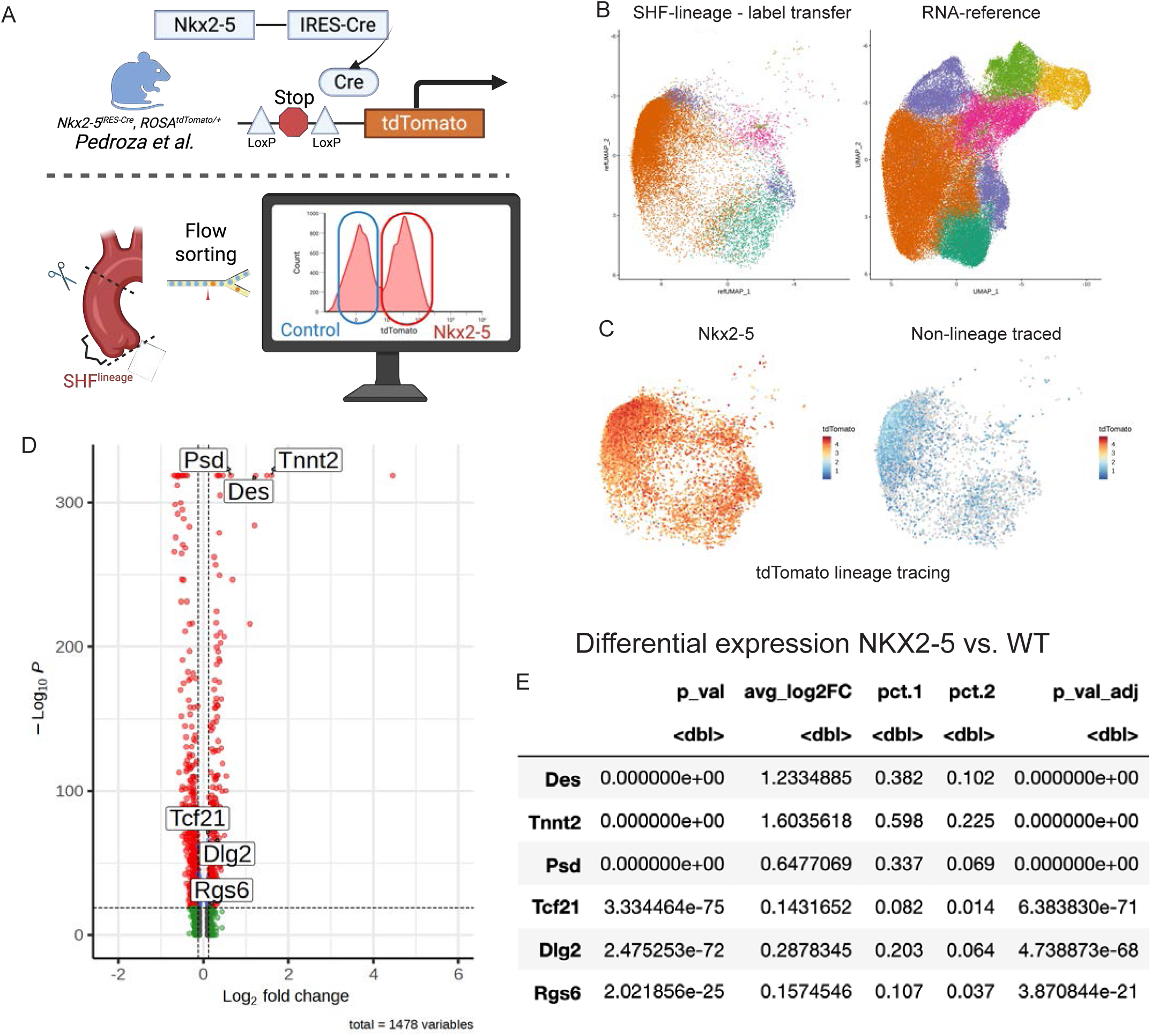
**A.** Schematic for *Nkx2-5* lineage tracing as described by Pedroza et al^21^. **B.** Cell projection and label transfer of *Nkx2-5* dataset onto timecourse control data. **C.** DimPlot for *Nkx2-5* dataset split by *Nkx2-5* lineage tracing status, showing *tdTomato* expression representing Nkx2-5 lineage traced cells. **D.** Volcano plot showing differential gene expression between *Nkx2-5* lineage traced cells vs non-lineage traced controls. **E.** Table of selected DEGs between Nkx2-5 and control datasets.

**Supp Fig 9.**
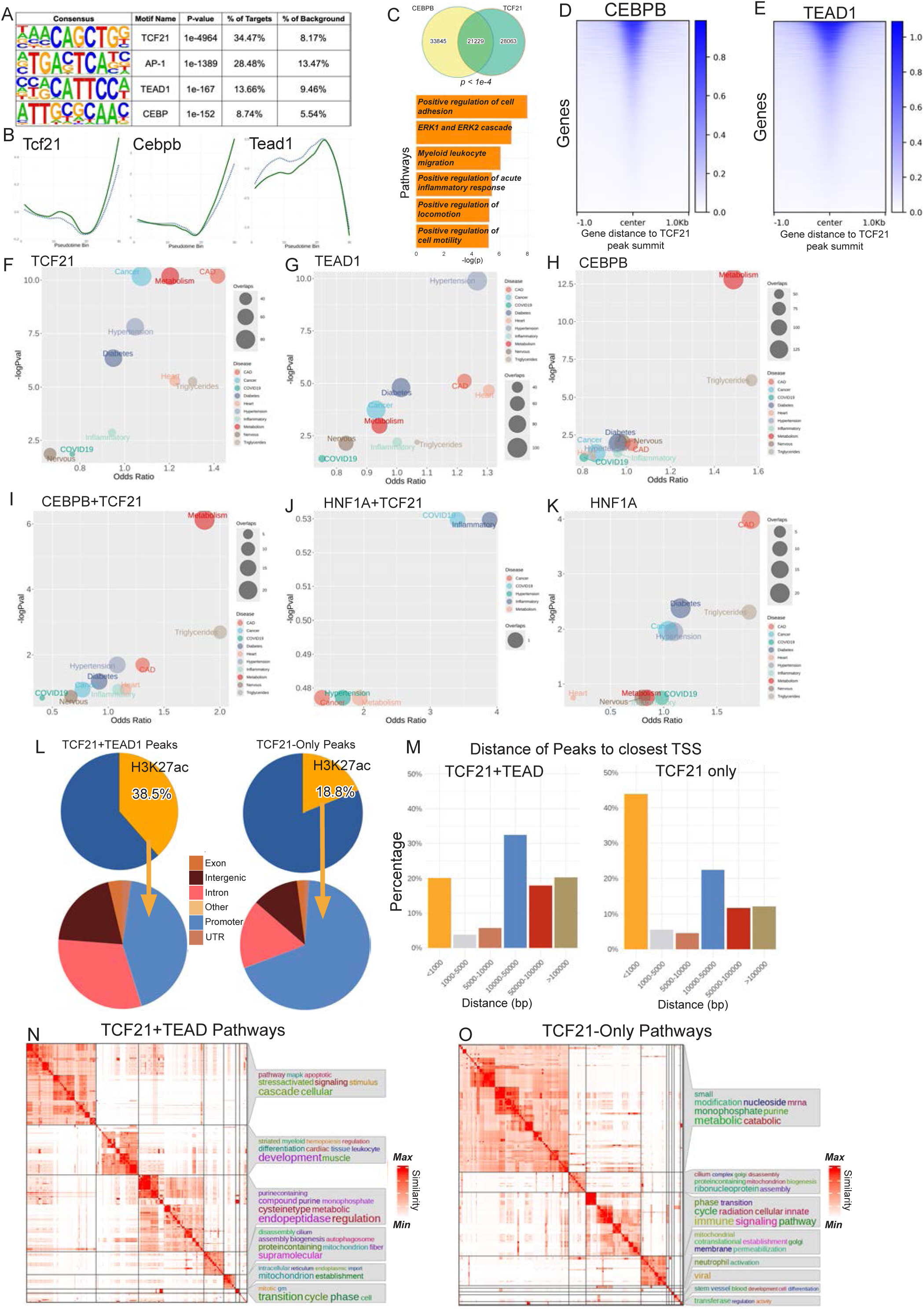
**A.** Top consensus TF motif sequences enriched amongst TCF21 HCASMC ChIPseq. **B.** Individual TF ChromVar scores for *Tcf21*, *Cebpb*, and *Tead1* across pseudotemporal bins comparing *Tcf21*- KO (dotted) and control (solid) SMC lineage cells. **C.** Overlapping ChIPseq peak number between *CEBPB* and *TCF21* and representative pathways identified by examining the intersected regions with the Genomic Regions Enrichment of Annotations Tool (GREAT). **D-E.** Heatmap of feature overlap for CEBPB+TCF21 and TEAD1+TCF21 ChIPseq **F-K.** GWASAnalytics results for GWAS SNP (EMBL-EBI GWAS Catalog) enrichment of overlapping ChIPseq peaks for TCF21, TEAD1, CEBPB, CEBPB+TCF21, HNF1A+TCF21 and HNF1A. HNF1A serves as negative control showing no significant CAD loci overlap with TCF21 despite both being CAD associated genes, suggesting of alternative mechanisms. **L.** ChIPseq annotation of TCF21+TEAD1 overlapping peaks and *TCF21*-only peaks. Top panel shows peak overlap with H3K27ac regions and bottom panel showing peak annotations. **M.** TSS distance analysis of TCF21+TEAD1 peaks and TCF21-only peaks. **N. O.** Representative pathways of overlapping TCF21+TEAD1 and TCF21-only ChIPseq peaks identified by GREAT.

## Materials and Methods

### Mouse strains, induction of lineage marker, and sample collection

Control (final genotype - *Tg^Myh^*^11^*^-CreERT^*^2^*, Tcf21^+/+^, ROSA^tdT/+^, ApoE^−/−^)* and *Tcf21* (final genotype - *Tg^Myh^*^11^*^-CreERT^*^2^*, Tcf21*^Δ*SMC/*Δ*SMC*^, *ROSA^tdT/+^, ApoE^−/−^)* knockout mice with floxed tdTomato fluorescent reporter to allow for SMC-specific lineage tracing were bred onto a C57BL/6 ApoE-/- background as previously described^16^. For all subsequent lineage tracing experiments, two doses of tamoxifen 48 hours apart via gavage was carried out when the mice reached 7.5 weeks of age. In the *Tcf21*-flox group, tamoxifen gavage also induces the *Tcf21* knockout in addition to tdTomato lineage tracing. Following tamoxifen treatment, all mice were placed on a high-fat diet. The study involved two primary high fat diet experimental arms: one for single-cell RNA sequencing (scRNA) and the other for Assay for Transposase-Accessible Chromatin sequencing (ATAC).

For scRNA analysis, samples were collected from Control mice at multiple time points named by number of weeks on high fat diet: 0, 3, 5, 7, 9, 12, and 16 weeks. Samples from the *Tcf21* knockout mice were collected at 5, 12, and 16 weeks. For ATAC analysis, samples were collected from the Control mice on the high-fat diet at 0, 5, 7, 9, 12, and 16 weeks and for *Tcf21* knockout mice at 5, 12, and 16 weeks.

### Aortic digestion for 10x Genomics microfluidics

Samples from mouse aorta were dissociated into single cells for single-cell RNA and ATAC sequencing using the 10x Genomics Chromium platform. Euthanized mice are perfused with phosphate buffered saline and dissected to obtain the aortic root up to the level of the brachiocephalic artery. The tissue is washed with PBS and incubated in an enzymatic dissociation cocktail containing Liberase (5401127001; Sigma-Aldrich) and Elastase (LS002279; Worthington) in Hank’s Balanced Salt Solution with calcium (HBSS+ Ca^2+^) for 30 minutes. Mechanical dissociation is performed for 5 minutes followed by visualization under the microscope to ensure single cell suspension. This suspension is strained through a 35µm nylon mesh snap cap into falcon test tubes (352235 BD) followed by FACS sorting (Sony SH800) for tdTomato positive and negative cells in parallel. For single cell ATAC, nuclei was isolated per 10X recommended protocol and captured on the 10X scATAC platform. Each individually sorted cell suspension was loaded onto 10x GEM G/H chips with remainder per 10x protocols (Chromium Single Cell 3’ RNA V3.1, Chromium Single Cell ATAC V2). For both scRNA and scATAC runs, samples at intermediate time points Weeks 7 and 9 were processed as pooled samples with tdTomato and non-tdTomato loaded at 1:1 ratio after FACS. Libraries were sequenced on the Illumina NovaSeq6000 platform with targeted depth of 40-50,000 reads per cell for RNA and 75,000 reads/cell for ATAC.

### Computational Methods

#### 10x RNA and ATAC Data Preprocessing

RNA fastq files were processed using Cellranger v7.0.1 (10x Genomics) to obtain transcript count matrices and aligned to mouse transcriptome mm10-2020-A-2.0.0 (10x Genomics) with custom addition of lineage tracing tdTomato transcript. Samples at each timepoint were aggregated and analyzed with R package with Seurat (v4.3)^60^. Low-quality cells and mitochondrial-rich cells were filtered with parameters mitochondrial percentage < 6%, ribosomal percentage < 25%, and nFeature_RNA > 1250 and < 8000. Gene expression count matrices underwent log-transformation and library-specific scaling. Importantly, no additional batch correction for visualization was required in the pre-processing steps given the uniform processing of mice samples. Principal component analysis was used for dimensionality reduction followed by clustering using the Louvain algorithm.

For the full timecourse dataset, aligned scRNA files from Control and *Tcf21*-KO data were merged and processed with identical QC parameters as above. Clustering was then performed on this merged dataset in order to ensure comparable differential gene analysis and pseudotime comparisons. The processed dataset is then split into Control and Tcf21-KO objects for independent analysis. We further applied logistic regression to determine the optimal *tdTomato* expression cutoff and subset on SMC-derived lineage cells exclusive to *tdTomato* fluorescence-activated cell sorting (FACS).

For force directed layouts, we followed methods as described in Schiebinger et al. 50 dimensional diffusion components are calculated with SCANPY (1.9.3) with default parameters. For each cell, its 20 nearest neighbors was used to produce a nearest neighbor map. Then we applied leiden clustering at a resolution of 0.36 and generated cluster connectivities via PAGA and visualized the force-directed layout on the k-NN graph using ForceAtlas2.

scATAC Raw fastq files were uniformly processed using Cellranger-atac-2.1.0 (10x Genomics) and aligned to cellranger-arc-mm10-2020-A-2.0.0 reference genome (10x Genomics). The data were processed to remove low-quality cells and reads with low mapping quality with parameters peak_region_fragments > 2500, peak_region_fragments < 100000, pct_reads_in_peaks > 30, nucleosome_signal <= 4, TSS.enrichment > 2. The remaining reads were then used to construct a sparse binary matrix, representing chromatin accessibility states across individual cells and genomic regions. A unified set of peaks from all samples (Control and *Tcf21*- KO) were merged using overlapping and adjacent peaks using CellRanger-ATAC aggr protocol. This unified peak set was re-quantified by term frequency-inverse document frequency (TF-IDF) using Signac^61^ (1.10) RunTFIDF().

To preprocess scRNA and scATAC data for integration, gene activities are first calculated from chromatin accessibility data using GeneActivity() function from Signac with default parameters and log normalization. Datasets were integrated using canonical correlation analysis (CCA) with the Seurat RunCCA() function upon 2000 features using the SelectIntegrationFeatures() in Seurat. We then use the integrated dataset to perform pseudotime ordering using Slingshot^62^ (2.6) and split the dataset into 30 pseudotime bins for uniform downstream analyses.

#### scATAC Functional and Motif Analyses

To generate identify overrepresented TF binding motifs, we performed hypergeometric enrichment on the top 20,000 variable peaks with GC-matched controls using the FindMotifs function in Signac. We then further merged this motif set with HOMER’s (4.10) de novo motif enrichment method on a per-cluster basis taking the top overrepresented motifs along with all similar motifs above a similarity score of 0.7. This allowed identification of a comprehensive set of enriched transcription factor motifs to aid summarization of ATAC data. For mouse ATAC data, the background peak set utilizing the entire merged peak set generated by CellRanger-ATAC (version 2.0.1) Aggr.

We then calculate a cluster specificity score by dividing the detection percentage of accessible chromatin for each cluster by the detection percentage of all other clusters with a minimum 10% cutoff for within cluster accessibility. We filtered peaks with a cluster specificity of >1.5 for FMC-1, FMC-2, and CMC and specificity > 1.25 for SMCs. From these peaks, we selected the top 5,000 peaks with the highest specificity score in the FMC-1, FMC-2, and CMC clusters while for the aggregated SMC cluster (SMC-1, SMC-2, SMC-3), we use the top 2500 peaks. ChromVAR^63^ was used to calculate the transcription factor specific enrichment scores of accessible motifs across each pseudotime bin through the Signac wrapper RunChromVAR().

#### Trajectory Analysis with Waddington Optimal Transport and CellRank2

To model and infer cellular trajectories in our data, we employed the Waddington optimal transport algorithm^30^. This approach infers cell growth rates using single-cell gene expression to generate a transition probability distribution, enabling the identification of cellular transitions and differentiation paths.

We applied the default command line implementation of Waddington WOT as described in Schiebinger et al. with cell scores derived from updated KEGG cell cycle and Apoptosis gene signatures for estimation of cell growth and death. Growth rate tables are extracted and added to single cell metadata for additional analyses. We generated a ‘fate correlation by comparing WOT predicted cell transition probability for the FMC-1, FMC-2 and CMC fate with each *Tcf21* downstream TF network module score in the combined *Tcf21*-KO and control dataset. As negative control, we also correlated respective TF expression with fate probability to identify consistently more robust correlation with TF module scores compared to its central TF.

Orthogonal trajectory analyses was performed using CellRank2. We applied the RealTimeKernel using default settings to identify terminal cell states and calculate driver gene trends plotted across our imported Slingshot derived pseudotime.

#### Data integration, gene regulatory network generation and in silico transcription factor perturbation

We utilize the Pando^45^ (1.0.1) to generate gene regulatory networks and CellOracle^46^ (0.18.0) for in silico TF perturbation experiments. By combining Pando’s network construction capabilities with CellOracle’s in silico perturbation analysis, this approach enables a comprehensive exploration of gene regulatory dynamics in single-cell data. We divided the integrated dataset into PseudoEarly (pseudotime bins 3-24) and PseudoLate (pseudotime bins 25-30) for further analyses.

To generate GRNs from our scRNA and scATAC integrated dataset, we identified transcription factors and their respective binding sites in regions of accessible chromatin through motif scanning, and these transcription factor module candidate regions linked to genes by proximity through the PANDO framework. The bagging regression model was selected to infer the relationships between TF expression, binding-site accessibility and target gene expression to generate cell state specific networks.

We utilized the extended transcription factor motif database from the original Pando manuscript (including JASPAR2020^64^, CIS-BP^65^ as well as TFs without known motifs assigned by sequence homology) and further included all JASPAR2020 database mouse reference motifs for downstream motif scanning. Gene regulatory network inference was then performed using default PANDO parameters for selection of candidate regulatory regions from scATAC data, transcription factor motif scanning, selection of region-TF pairs, while the final regression model, we substitute the bagging ridge model for the default generalized linear model to match that of CellOracle. The output coefficient table was extracted and filtered for adjusted p-value < 0.05, R^2^ > 0.1, minimum number of variables (region-TF pairs) per model > 10, and minimum genes per module > 5 to generate the final regulatory graph that is provided as input for downstream analysis including:

1. Weighted pseudotime x TF module UMAP, is calculated by the get_network_graph() function in Pando to generate a subgraph with significant module TFs as features to generate a UMAP embedding which takes into account the ingoing connection of each node (TF) as well as coexpression of TFs. Additionally, the product of average pseudotime per cell by TF expression is colorized onto each TF module to provide a visual representation of the TF module relationships in the context of pseudotime.
2. TF activity score – calculated by the averaging the sum of TF expression x target gene coefficient for each TF module in the early (pseudotime bin 3-24) or late (pseudotime bin 25-30) SMC gene regulatory network.

For in silico perturbation, the merged control time course integrated dataset is downsampled to 40,000 cells to allow for optimal computation speed along with the top 3000 most variable genes calculated in Seurat and extended TF database from Pando converted into CellOracle compatible format to generate a baseline ‘Oracle’ file. The Pando derived TF-gene regulatory network table is converted into ‘links’ network file format as input for CellOracle’s link_data parameter for in silico systemic TF perturbation modeling with negative PS sums visualized as described in the CellOracle tutorial.

#### Mouse to human scRNA label transfer and Spatial Slideseq analysis

Data is obtained and re-processed from the CZI Arterial Atlas courtesy of Zhao et al. ^23^. We processed single cell RNA data based on parameters published in Zhao et al. and subset the SMC clusters. We then transferred labels from our control smooth muscle control dataset and transferred labels using CCA with 1:30 dim and assigned SMC cell states to the human scRNA data. Slide-Seq object was aligned to hg38 by curioseeker pipeline and again underwent processing as described in Zhao et al. ^23^. Robust Cell Type Decomposition was then applied to integrate this label transferred reference with spot-level data from the spatial transcriptomic sequencing using Seurat V5^66^. Visualization was performed using Seurat built-in functions including SpatialDimPlot to visualize the label transferred cell states.

#### ChIPseq analyses

Fastq files were mapped to hg38/GRCh37 genome with Bowtie2 (1.2.3). ChIP-seq peaks were then called with MACS2 (2.2.7.1) using default parameters. From this output, ‘robust’ peaks were selected by specifying a minimum fold-enrichment of 5.

#### GWAS trait SNP enrichment analyses

The intersection of GWAS loci and transcription factor binding (*TEAD1* + *TCF21*) was defined as the SNPs located within any overlapping region with ChIP-seq peaks. Direct overlap of SNPs assimilated from GWAS Catalog + MVP CAD GWAS was performed with GWASanalytics script (https://github.com/zhaoshuoxp/GWASanalytics). The binomial overlap performed by this script has been described previously ^67^.

#### scDRS (Single Cell Disease Relevance Score)

The scDRS^53^ algorithm calculates a disease relevance score for each cell by comparing its gene expression pattern with a reference disease gene signature obtained from the MVP coronary artery disease GWAS data. We utilized summary statistics obtained from the coronary artery disease GWAS meta-analysis from the Million Veterans Program (MVP) which also incorporates transancestry genetic data. MVP CAD GWAS was munged using (MAGMA) to obtained a weighted CAD-associated gene list that is applied towards the mouse timecourse data. Remainder scDRS analysis was performed as described in Zhang et al. ^53^.

#### Curated CAD GWAS gene list

A curated list of nominated GWAS genes based on prior CAD GWAS meta-analysis was tabulated from Erdmann 2018, Koyama 2019, Matsunaga 2020, Tcheandjieu 2022, Aragam 2022 ^5,6,68–70^

#### Heritability adjusted PrediXcan, LocusZoom and eQTL correlation visualization

We estimated gene expression risk directionality inferred from the updated heritability adjusted S-PrediXcan modeling ^54^ which was use. LocusZoomR^71^ (0.3.8) was used to visualize gene locus plots. eQTL colocalization was performed by extracting positional coordinates from MVP CAD GWAS and retrieving SNPs from dbGaP. SNP with lowest p-value near nominated GWAS gene was selected and all SNPs meeting GWAS significance 5e-8 and within 50kb up and downstream of lead SNP were selected. We then identify which SNPs had corresponding eQTLs using the V10 GTEx release. The beta for each SNP was correlated with the corresponding NES of each eQTL and plotted as a scatter graph with color representing the R^2^ (calculated with LDlinkR) with lead SNP.

### Experimental Methods

#### In situ RNA hybridization (RNAScope)

Slides were processed according to the manufacturer’s instructions, using reagents from ACD Bio (ACD 322360-USM). Slides were washed in PBS, then immersed in 1× Target Retrieval reagent at 100°C for 5 minutes. After washing twice in deionized water, slides were immersed in 100% ethanol, air-dried, and sections were encircled with a liquid-blocking pen. Sections were incubated with Protease III reagent at 40°C for 30 minutes, then washed twice with deionized water. Sections were incubated with probes against Fbln1 (ACD 502881), Fbln2 (ACD 447931), Ibsp (ACD 415501), C3 (ACD 417841), Tcf21 (ACD 508661), Thbs1 (ACD 457891), Notch3(ACD 425171), Col8a1 (ACD 518071), Ttc9 (ACD 1113921-C1), Loxl1 (ACD 492531), Bmp1 (ACD 311151), Vcam1 (ACD 438641), Col2a1 (ACD 407221) or a negative control probe (ACD 310043) for 3 hours at 40°C. Multiplex fluorescence and colorimetric assays were performed per the manufacturer’s instructions.

#### HCASMC and A7r5 cell culture

Primary HCASMCs and HCASMC-hTERT ^72^ were maintained in SMC basal medium (SmBM) supplemented with SmGM-2 SingleQuots kit (Lonza CC-3182) including human epidermal growth factor, insulin, human basic fibroblast growth factor and 5% FBS, according to the manufacturer’s instructions.

Rat aortic smooth muscle cells (A7r5) were purchased from ATCC and cultured in Dulbecco’s Modified Eagle Medium (DMEM) high glucose (Fisher Scientific, #MT10013CV) with 10% FBS at 37 °C and 5% CO2. A7r5 at passage 6-9 were used for experiments.

#### CEBPB, TEAD, H3K27ac ChIPseq

Approximately 1,000,000 HCASMCs were cross linked in 1% formaldehyde for 10 minutes and washed with PBS and replaced with hypotonic buffer (20 mM Hepes (pH 7.9), 10 mM KCl, 1 mM EDTA (pH 8) and 10% glycerol) and incubated on ice for 6 minutes. Cells were then sonicated using a Branson 250 Sonifier (using power setting 5, constant duty for 10 rounds of 30-second pulses) with confirmation of chromatin fragments at 250-400 babse pairs. Lysates were then incubated overnight with 5 µg of anti-CEBPB (Santa Cruz sc-150), TEAD(Cell Signaling 12292S), or H3K27ac(Abcam 4729). Protein-DNA complexes were captured with Protein G agarose beads (Sigma 8104) and eluted in 1% SDS TE buffer at 65 °C. After reverse cross-linking followed by RNase A and proteinase K digestion, chromatin was purified using a QIAquick PCR purification kit (Qiagen 28706). ChIP DNA sequencing libraries were prepared using the Kappa HyperPrep (Roche 07962347001) and sequenced on a NovaSeq6000 with 150-base pair paired-end reads.

#### Proximity Ligation Assay

PLA was performed using manufacturer provided protocols for Sigma DuoLink In Situ Red Start Kit Mouse/Rabbit (DUO92101) with antibodies to TCF21 (Sigma AV33421), TEAD1 (sc-376113) and anti-mouse IgG (Vector I-2000-1).

#### Dual luciferase assays

Enhancer/Promoter elements (SRF 2nd intron; *chr15:73633709-73634109*, BMP1 6th intron; *chr8:22178981- 22179700*, Loxl1 TSS; *chr15:73926050-73926450*) demonstrating overlapping between *Tcf21* and *Tead* ChIP-seq sites were cloned into pWPI and transfected into A7r5 cells. A7r5 cells were seeded into 24 well plate (1.5 × 104 cells/well) in DMEM containing 10% FBS and incubated at 37 °C and 5% CO2 overnight. Cells were transfected with varying combinations of luciferase reporter plasmids (pLuc-MCS (empty), pLuc-enhancer, cDNAs (pWPI (empty), pWPI-TCF21, pWPI-TEAD1 and pWPI-MYOCD), and Renilla luciferase plasmid using Lipofectamine 3000 (Invitrogen, #L3000015). Six hours after transfection, the media was changed to fresh complete media. Relative luciferase activity (firefly/Renilla luciferase ratio) was measured by SpectraMax L luminometer (Molecular Devices) 24 h after transfection. All experiments were conducted in at least triplicate and normalized to the reporter plasmid after subtracting empty luciferase construct luminescence.

### Statistical Methods

To identify differentially expressed genes in the scRNA-Seq data, we split our data by PseudoEarly (pseudotime bins 3-24) and PseudoLate (pseudotime bins 25-30) and employed the FindMarkers function with the “DESeq2” wrapper and filter for genes with absolute value Log2FC > 0.15. For differential scATAC peak analysis, we applied the likelihood ratio test ‘LR’ from FindMarkers and used nCount_peaks as the latent.variable and min.pct as 0.001 on a per cell type cluster basis. For differential ChromVar analysis, we calculated row means and calculated the average difference in Z-score between conditions using the wilcox test from FindMarkers.

For Jensen-shannon divergence (JSD) calculation we compared the ChromVar motif deviation distribution for each motif across pseudotime by calculating the Kullback-Leibler (KL) divergence from each distribution to the midpoint distribution where:

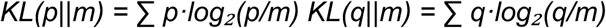

Where p and q represent the two distributions being compared and m is the midpoint distribution calculated as the average of two normalized distributions

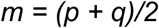

Then the JSD was computed as the square root of the average of the two KL divergences.

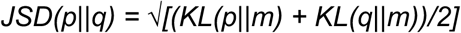

We first randomly split the control dataset equally and calculated the JSD divergence to derive a control distribution for each motif. We then calculated JSD divergence between Tcf21-KO and control.

To determine statistical significance, we constructed a null distribution using the control JSD measurements by performing 10,000 random sampling with replacement from the control values. One-sided empirical p-values were calculated for each TF using the formula p = (number of permuted values > observed value +1) / (total number of permutations + 1).

Fisher’s exact test was used to assess the significance of overlaps between genomic regions. Adjusted P-values are corrected with Benjamini-Hochberg procedure and <0.05 were considered statistically significant. All error bars represent standard error of the mean (SE).

## Data and code availability

All single cell data are deposited to CellxGene to be made available for public access.

All ChIPseq data generated and analyzed (CEBPB, H3K27ac, TEAD1) are deposited to National Center for Biotechnology Information Gene Expression Omnibus (GEO) to be made available for public access.

TCF21-pooled ChIPseq and HNF1A ChIPseq are downloaded from (GEO): GSE141752 GWAS Catalog data were downloaded from https://www.ebi.ac.uk/gwas/ and Million Veteran Program (MVP) were downloaded from dbGap with accession number phs001672.v3.p1

## Acknowledgements

This work was supported by National Institutes of Health grants R01HL171045 (TQ), R01HL134817 (TQ), R01HL139478 (TQ), R01HL156846 (TQ), R01HL158525 (TQ), UM1HG011972 (TQ), U01HG011762 (TQ), and the William G. Irwin Foundation (TQ), as well as a Human Cell Atlas grant (ZF2019-002437) from the Chan Zuckerberg Foundation (TQ). Support was provided to DL through the NIH grant F32HL165819.

## Author Contributions

T.Q., R.W, and A. K, conceived and supervised the research plan. R. W, D. L., P.C., S. K., J.M., W.G., W.J., S.D., B.P., C.W., performed single cell captures and single cell analyses, D. L., and T. N. performed experiments with cultured cells, and helped with genomic analyses. D.L., R.K, R.W., P.C., W.J., maintained mouse colonies and performed RNAScope experiments, D. L., S. K., R. W. and Q.Z. performed analyses. D. L., and T. Q. wrote the manuscript, R.W., S. K. contributed to writing and proof reading.

## References

1 Roth, G. A. et al. Global Burden of Cardiovascular Diseases and Risk Factors, 1990-2019: Update From the GBD 2019 Study. J Am Coll Cardiol 76, 2982–3021 (2020). 10.1016/j.jacc.2020.11.010

2 Khera, A. V. & Kathiresan, S. Genetics of coronary artery disease: discovery, biology and clinical translation. Nat Rev Genet 18, 331–344 (2017). 10.1038/nrg.2016.160

3 Zdravkovic, S. et al. Heritability of death from coronary heart disease: a 36-year follow-up of 20 966 Swedish twins. J Intern Med 252, 247–254 (2002).

4 Marenberg, M. E., Risch, N., Berkman, L. F., Floderus, B. & de Faire, U. Genetic susceptibility to death from coronary heart disease in a study of twins. N Engl J Med 330, 1041–1046 (1994). 10.1056/NEJM199404143301503

5 Tcheandjieu, C. et al. Large-scale genome-wide association study of coronary artery disease in genetically diverse populations. Nat Med 28, 1679–1692 (2022). 10.1038/s41591-022-01891-3

6 Aragam, K. G. et al. Discovery and systematic characterization of risk variants and genes for coronary artery disease in over a million participants. Nature genetics 54, 1803–1815 (2022). 10.1038/s41588-022-01233-6

7 Quertermous, T. et al. Genome-Wide Genetic Associations Prioritize Evaluation of Causal Mechanisms of Atherosclerotic Disease Risk. Arterioscler Thromb Vasc Biol 44, 323–327 (2024). 10.1161/ATVBAHA.123.319480

8 Turner, A. W. et al. Single-nucleus chromatin accessibility profiling highlights regulatory mechanisms of coronary artery disease risk. Nature genetics 54, 804–816 (2022). 10.1038/s41588-022-01069-0

9 Ord, T. et al. Dissecting the polygenic basis of atherosclerosis via disease-associated cell state signatures. American journal of human genetics 110, 722–740 (2023). 10.1016/j.ajhg.2023.03.013

10 Zhang, K. et al. A single-cell atlas of chromatin accessibility in the human genome. Cell 184, 5985–6001 e5919 (2021). 10.1016/j.cell.2021.10.024

11 Alencar, G. F. et al. The Stem Cell Pluripotency Genes Klf4 and Oct4 Regulate Complex SMC Phenotypic Changes Critical in Late-Stage Atherosclerotic Lesion Pathogenesis. Circulation 142, 2045–2059 (2020).

12 Cheng, P. et al. Smad3 regulates smooth muscle cell fate and mediates adverse remodelling and calcification of the atherosclerotic plaque. Nature Cardiovascular Research 4, 322–333 (2022).

13 Cheng, P. et al. ZEB2 Shapes the Epigenetic Landscape of Atherosclerosis. Circulation (2022). 10.1161/CIRCULATIONAHA.121.057789

14 Kim, J. B. et al. Environment-Sensing Aryl Hydrocarbon Receptor Inhibits the Chondrogenic Fate of Modulated Smooth Muscle Cells in Atherosclerotic Lesions. Circulation 142, 575–590 (2020). 10.1161/CIRCULATIONAHA.120.045981

15 Pan, H. et al. Single-Cell Genomics Reveals a Novel Cell State During Smooth Muscle Cell Phenotypic Switching and Potential Therapeutic Targets for Atherosclerosis in Mouse and Human. Circulation (2020). 10.1161/CIRCULATIONAHA.120.048378

16 Wirka, R. et al. Single cell analysis of smooth muscle cell phenotypic modulation in vivo reveals a critical role for coronary disease gene TCF21 in mice and humans. Nat Med 25, 1280–1289 (2019).

17 Kim, H. J. et al. Molecular mechanisms of coronary artery disease risk at the PDGFD locus. Nat Commun 14, 847 (2023). 10.1038/s41467-023-36518-9

18 Shao, X. et al. Integrated single-cell RNA-seq analysis reveals the vital cell types and dynamic development signature of atherosclerosis. Front Physiol 14, 1118239 (2023). 10.3389/fphys.2023.1118239

19 Sharma, D. et al. Comprehensive Integration of Multiple Single-Cell Transcriptomic Data Sets Defines Distinct Cell Populations and Their Phenotypic Changes in Murine Atherosclerosis. Arterioscler Thromb Vasc Biol 44, 391–408 (2024). 10.1161/ATVBAHA.123.320030

20 Lin, C. J. et al. Distinct Patterns of Smooth Muscle Phenotypic Modulation in Thoracic and Abdominal Aortic Aneurysms. J Cardiovasc Dev Dis 11 (2024). 10.3390/jcdd11110349

21 Pedroza, A. J. et al. Embryologic Origin Influences Smooth Muscle Cell Phenotypic Modulation Signatures in Murine Marfan Syndrome Aortic Aneurysm. Arterioscler Thromb Vasc Biol 42, 1154–1168 (2022). 10.1161/ATVBAHA.122.317381

22 Shukla, S. et al. Single-Cell Transcriptomics Identifies Selective Lineage-Specific Regulation of Genes in Aortic Smooth Muscle Cells in Mice. Arterioscler Thromb Vasc Biol (2025). 10.1161/ATVBAHA.124.321482

23 Zhao, Q. et al. A cell and transcriptome atlas of the human arterial vasculature. bioRxiv (2024). 10.1101/2024.09.10.612293

24 Acharya, A. et al. The bHLH transcription factor Tcf21 is required for lineage-specific EMT of cardiac fibroblast progenitors. Development 139, 2139–2149 (2012). 10.1242/dev.079970

25 Misra, A. et al. Integrin beta3 regulates clonality and fate of smooth muscle-derived atherosclerotic plaque cells. Nat Commun 9, 2073 (2018). 10.1038/s41467-018-04447-7

26 Worssam, M. D. et al. Cellular mechanisms of oligoclonal vascular smooth muscle cell expansion in cardiovascular disease. Cardiovascular research 119, 1279–1294 (2023). 10.1093/cvr/cvac138

27 Jacobsen, K. et al. Diverse cellular architecture of atherosclerotic plaque derives from clonal expansion of a few medial SMCs. JCI Insight 2 (2017). 10.1172/jci.insight.95890

28 Chappell, J. et al. Extensive Proliferation of a Subset of Differentiated, yet Plastic, Medial Vascular Smooth Muscle Cells Contributes to Neointimal Formation in Mouse Injury and Atherosclerosis Models. Circulation research 119, 1313–1323 (2016). 10.1161/CIRCRESAHA.116.309799

29 Haseeb, A. et al. SOX9 keeps growth plates and articular cartilage healthy by inhibiting chondrocyte dedifferentiation/osteoblastic redifferentiation. Proceedings of the National Academy of Sciences of the United States of America 118 (2021). 10.1073/pnas.2019152118

30 Schiebinger, G. et al. Optimal-Transport Analysis of Single-Cell Gene Expression Identifies Developmental Trajectories in Reprogramming. Cell 176, 928–943 e922 (2019). 10.1016/j.cell.2019.01.006

31 Zhang, S., Afanassiev, A., Greenstreet, L., Matsumoto, T. & Schiebinger, G. Optimal transport analysis reveals trajectories in steady-state systems. PLoS Comput Biol 17, e1009466 (2021). 10.1371/journal.pcbi.1009466

32 Mosquera, J. V. et al. Integrative single-cell meta-analysis reveals disease-relevant vascular cell states and markers in human atherosclerosis. Cell Rep 42, 113380 (2023). 10.1016/j.celrep.2023.113380

33 Witzenbichler, B. et al. Regulation of smooth muscle cell migration and integrin expression by the Gax transcription factor. The Journal of clinical investigation 104, 1469–1480 (1999). 10.1172/JCI7251

34 Jeon, B. N. et al. KR-POK interacts with p53 and represses its ability to activate transcription of p21WAF1/CDKN1A. Cancer research 72, 1137–1148 (2012). 10.1158/0008-5472.CAN-11-2433

35 Tanaka, T. et al. Runx2 represses myocardin-mediated differentiation and facilitates osteogenic conversion of vascular smooth muscle cells. Molecular and cellular biology 28, 1147–1160 (2008). 10.1128/MCB.01771-07

36 Nagao, M. et al. Coronary Disease Associated Gene TCF21 Inhibits Smooth Muscle Cell Differentiation by Blocking the Myocardin-Serum Response Factor Pathway. Circulation research 126, 517–529. (2019). 10.1161/CIRCRESAHA.119.315968

37 Bonnet, S. et al. The nuclear factor of activated T cells in pulmonary arterial hypertension can be therapeutically targeted. Proceedings of the National Academy of Sciences of the United States of America 104, 11418–11423 (2007). 10.1073/pnas.0610467104

38 Li, M. et al. Sildenafil inhibits calcineurin/NFATc2-mediated cyclin A expression in pulmonary artery smooth muscle cells. Life Sci 89, 644–649 (2011). 10.1016/j.lfs.2011.07.023

39 Canalis, E., Schilling, L., Eller, T. & Yu, J. Role of nuclear factor of activated T cells in chondrogenesis osteogenesis and osteochondroma formation. J Endocrinol Invest 45, 1507–1520 (2022). 10.1007/s40618-022-01781-y

40 Engleka, K. A. et al. Islet1 derivatives in the heart are of both neural crest and second heart field origin. Circulation research 110, 922–926 (2012). 10.1161/CIRCRESAHA.112.266510

41 Liu, C. F., Samsa, W. E., Zhou, G. & Lefebvre, V. Transcriptional control of chondrocyte specification and differentiation. Semin Cell Dev Biol 62, 34–49 (2017). 10.1016/j.semcdb.2016.10.004

42 Mackie, E. J., Ahmed, Y. A., Tatarczuch, L., Chen, K. S. & Mirams, M. Endochondral ossification: how cartilage is converted into bone in the developing skeleton. Int J Biochem Cell Biol 40, 46–62 (2008). 10.1016/j.biocel.2007.06.009

43 McLean, C. Y. et al. GREAT improves functional interpretation of cis-regulatory regions. Nature biotechnology 28, 495–501 (2010). 10.1038/nbt.1630

44 Bennett, M. R., Sinha, S. & Owens, G. K. Vascular Smooth Muscle Cells in Atherosclerosis. Circulation research 118, 692–702 (2016). 10.1161/CIRCRESAHA.115.306361

45 Fleck, J. S. et al. Inferring and perturbing cell fate regulomes in human brain organoids. Nature (2022). 10.1038/s41586-022-05279-8

46 Kamimoto, K. et al. Dissecting cell identity via network inference and in silico gene perturbation. Nature 614, 742–751 (2023). 10.1038/s41586-022-05688-9

47 Lee, S. Y. et al. Differential but complementary roles of HIF-1alpha and HIF-2alpha in the regulation of bone homeostasis. Commun Biol 7, 892 (2024). 10.1038/s42003-024-06581-z

48 Salminen, A. Mutual antagonism between aryl hydrocarbon receptor and hypoxia-inducible factor- 1alpha (AhR/HIF-1alpha) signaling: Impact on the aging process. Cellular signalling 99, 110445 (2022). 10.1016/j.cellsig.2022.110445

49 Lambert, J. et al. Network-based prioritization and validation of regulators of vascular smooth muscle cell proliferation in disease. Nat Cardiovasc Res 3, 714–733 (2024). 10.1038/s44161-024-00474-4

50 Lin, M. E. et al. Runx2 deletion in smooth muscle cells inhibits vascular osteochondrogenesis and calcification but not atherosclerotic lesion formation. Cardiovasc Res 112, 606–616 (2016). 10.1093/cvr/cvw205

51 Zhao, Q. et al. TCF21 and AP-1 interact through epigenetic modifications to regulate coronary artery disease gene expression. Genome Med 11, 23 (2019).

52 Zhao, Q. et al. Molecular mechanisms of coronary disease revealed using quantitative trait loci for TCF21 binding, chromatin accessibility, and chromosomal looping. Genome Biol 21, 135 (2020). 10.1186/s13059-020-02049-5

53 Zhang, M. J. et al. Polygenic enrichment distinguishes disease associations of individual cells in single-cell RNA-seq data. Nature genetics 54, 1572–1580 (2022). 10.1038/s41588-022-01167-z

54 Liang, Y., Nyasimi, F. & Im, H. K. Pervasive polygenicity of complex traits inflates false positive rates in transcriptome-wide association studies. bioRxiv (2024). 10.1101/2023.10.17.562831

55 Huang, C. K. et al. Androgen receptor promotes abdominal aortic aneurysm development via modulating inflammatory interleukin-1alpha and transforming growth factor-beta1 expression. Hypertension 66, 881–891 (2015). 10.1161/HYPERTENSIONAHA.115.05654

56 Sun, Y. et al. Smooth muscle cell-specific runx2 deficiency inhibits vascular calcification. Circulation research 111, 543–552 (2012). 10.1161/CIRCRESAHA.112.267237

57 Topouzis, S. & Majesky, M. W. Smooth muscle lineage diversity in the chick embryo. Two types of aortic smooth muscle cell differ in growth and receptor-mediated transcriptional responses to transforming growth factor-beta. Developmental biology 178, 430–445 (1996). 10.1006/dbio.1996.0229

58 Madura, J. A., 2nd et al. Regional differences in platelet-derived growth factor production by the canine aorta. J Vasc Res 33, 53–61 (1996). 10.1159/000159132

59 Trigueros-Motos, L. et al. Embryological-origin-dependent differences in homeobox expression in adult aorta: role in regional phenotypic variability and regulation of NF-kappaB activity. Arterioscler Thromb Vasc Biol 33, 1248–1256 (2013). 10.1161/ATVBAHA.112.300539

60 Hao, Y. et al. Integrated analysis of multimodal single-cell data. Cell 184, 3573–3587 e3529 (2021). 10.1016/j.cell.2021.04.048

61 Stuart, T., Srivastava, A., Madad, S., Lareau, C. A. & Satija, R. Single-cell chromatin state analysis with Signac. Nature methods 18, 1333–1341 (2021). 10.1038/s41592-021-01282-5

62 Street, K. et al. Slingshot: cell lineage and pseudotime inference for single-cell transcriptomics. BMC Genomics 19, 477 (2018). 10.1186/s12864-018-4772-0

63 Schep, A. N., Wu, B., Buenrostro, J. D. & Greenleaf, W. J. chromVAR: inferring transcription-factor-associated accessibility from single-cell epigenomic data. Nature methods 14, 975–978 (2017). 10.1038/nmeth.4401

64 Fornes, O. et al. JASPAR 2020: update of the open-access database of transcription factor binding profiles. Nucleic acids research 48, D87–D92 (2020). 10.1093/nar/gkz1001

65 Weirauch, M. T. et al. Determination and inference of eukaryotic transcription factor sequence specificity. Cell 158, 1431–1443 (2014). 10.1016/j.cell.2014.08.009

66 Hao, Y. et al. Dictionary learning for integrative, multimodal and scalable single-cell analysis. Nature biotechnology 42, 293–304 (2024). 10.1038/s41587-023-01767-y

67 Kim, J. B. et al. TCF21 and the environmental sensor aryl-hydrocarbon receptor cooperate to activate a pro-inflammatory gene expression program in coronary artery smooth muscle cells. PLoS Genet 13, e1006750 (2017). 10.1371/journal.pgen.1006750

68 Koyama, S. et al. Population-specific and transethnic genome-wide analyses reveal distinct and shared genetic risks of coronary artery disease. bioRxiv, 827550 (2019). 10.1101/827550

69 Matsunaga, H. et al. Transethnic Meta-Analysis of Genome-Wide Association Studies Identifies Three New Loci and Characterizes Population-Specific Differences for Coronary Artery Disease. Circ Genom Precis Med 13, e002670 (2020). 10.1161/CIRCGEN.119.002670

70 Erdmann, J., Kessler, T., Munoz Venegas, L. & Schunkert, H. A decade of genome-wide association studies for coronary artery disease: the challenges ahead. Cardiovascular research 114, 1241–1257 (2018). 10.1093/cvr/cvy084

71 Lewis, M. J. & Wang, S. locuszoomr: an R package for visualizing publication-ready regional gene locus plots. Bioinform Adv 5, vbaf006 (2025). 10.1093/bioadv/vbaf006

72 Kavousi, M. et al. Multi-ancestry genome-wide study identifies effector genes and druggable pathways for coronary artery calcification. Nature genetics 55, 1651–1664 (2023). 10.1038/s41588-023-01518-4

